# Locus coeruleus activation transforms cortical taste representations

**DOI:** 10.64898/2026.06.02.729661

**Authors:** Will Fan, Paula To, Natale R. Sciolino

## Abstract

Norepinephrine neurons in the locus coeruleus (LC) shape sensory responses across the brain, yet how LC activity reorganizes population representations of behaviorally relevant sensory attributes remains unclear. We addressed this question in the primary gustatory cortex (GC), a system largely unexplored in neuromodulation research and well-suited for linking sensory coding to affective value. Using miniscope calcium imaging in awake mice combined with optogenetic LC activation, we tested how phasic and tonic LC activity modulate GC encoding of three taste attributes: palatability, mixture ratio, and intensity. Phasic LC activation enhanced correlations between neuronal responses and tastant palatability and expanded the dynamic range of taste representations along a palatability-relevant axis. This expansion was driven primarily by an aversive shift in the representation of all tastants except sucrose, the most palatable stimulus. For mixture ratio and concentration, LC activation induced both stretching and rotation of attribute axes, potentially reflecting dependencies between these attributes and palatability. Analysis of LC-induced changes in neuronal tuning revealed that these population-level transformations likely arise from a combination of multiplicative gain modulation and flexible, tastant-specific changes in tuning. In contrast, tonic LC activation engaged fewer GC neurons and did not produce stretching along any attribute axis, demonstrating that LC effects on population coding depend on activation pattern. Together, these findings provide causal evidence that phasic LC activation reshapes GC population geometry to preferentially enhance palatability encoding, establishing a framework for understanding how neuromodulation may bias taste representations relevant to feeding behavior.

## INTRODUCTION

Sensory processing is adaptively shaped by an animal’s internal state, including fluctuations in arousal, attention, motivation, and stress^1–6^. Among the neuromodulatory systems that mediate these state-dependent changes, the noradrenergic neurons of the locus coeruleus (LC) are one of the most prominent and extensively studied sources of modulation in sensory pathways^7–10^. LC activity closely tracks behavioral state and is commonly described in terms of phasic and tonic firing modes^11^: phasic activation is linked to vigilance, attentional shifts, and motivated actions^12–14^, whereas elevated tonic firing predominates during stress and hyperarousal^15,16^. Through its widespread projections to nearly all sensory networks^17^, the LC can broadly influence circuit dynamics through norepinephrine (NE) release. LC stimulation and pharmacological NE application produce diverse effects across sensory regions, including bidirectional modulation of responses^18–20^, changes in tuning selectivity^21–23^, improved signal-to-noise ratio^24,25^, and alterations in response timing^18,19,26^. Additionally, LC activation shapes sensory processing in a manner dependent on brain region^18,27^ and LC activity pattern^26,28^. However, because these effects have largely been characterized at the level of individual neurons, it remains unclear how LC activity reorganizes sensory representations at the level of neural populations—the scale at which perception and behavior ultimately emerge.

The advantage of moving beyond single-neuron analyses to population-level frameworks^29–31^ is that the latter more directly link neural activity to perception, action, and cognition^32,33^. In sensory systems, the instantaneous population response to a stimulus can be represented as a point in high-dimensional activity space, where each axis corresponds to the activity of one neuron. Across stimuli, these responses patterns form a structured, low-dimensional manifold that captures the relationships among stimuli as encoded by the population^34^. Within this geometric framework, neuromodulation corresponds to transformations of the coding space—scaling, shifting, rotating, or warping the representational manifold. Different forms of geometric transformations make distinct predictions about feature encoding, discriminability, and downstream readout. Despite the promise of this approach, only a small number of studies have applied some population-based analyses to LC modulation^27,28,35^, and how LC activity casually transforms sensory coding geometry remains poorly understood. Determining how LC activity reshapes representational geometry could reveal general principals by which neuromodulators bias sensory processing toward behaviorally relevant stimulus attributes.

In the present study, we investigated how distinct patterns of LC activation influence taste representations in the primary gustatory cortex (GC), examining effects at both the single-neuron and population levels. The GC is a powerful model system for several reasons. First, compared to other sensory modalities, gustation remains underrepresented in the neuromodulation literature, with no prior work directly testing the causal impact of LC activity on GC coding. Second, GC population activity encodes not only taste identity and intensity, but also *palatability*^36–39^—a behaviorally relevant dimension that links sensory coding to affective processing and feeding-related behavior. Third, palatability coding in GC is flexibly regulated by behavioral states such as attention^40^ and sickness^41^, but the pathways mediating these effects remain unknown. Given LC’s role in adaptive sensory-guided decisions^27,42–44^, we hypothesized that LC activation would reorganize GC population geometry by enhancing palatability representations, among other impacts.

To test this hypothesis, we combined miniscope calcium imaging in GC with optogenetic LC activation in awake mice receiving passively delivered taste stimuli. We examined the impact of phasic and tonic LC activation on three taste attributes: palatability, mixture ratio, and concentration. Phasic LC activation enhanced the correlation between neuronal responses and tastant palatability and stretched the dynamic range of basic taste representations along a palatability-relevant axis. Similar stretching occurred for stimuli varying in mixture ratio and intensity; however, the population axes relevant to these dimensions underwent rotations, potentially due to their partial correlation with palatability. These population-level transformations likely arise from combined effects of gain modulation and flexible tuning changes at the single-neuron level. In contrast to phasic stimulation, prolonged tonic stimulation—despite inducing greater NE release—modulated a smaller fraction of GC neurons and produced no stretching along any attribute axis. Taken together, our findings demonstrate that LC activity shapes GC population coding in a pattern-dependent manner, establishing a causal framework for understanding how neuromodulation reorganizes cortical sensory representations of behaviorally relevant stimulus attributes.

## RESULTS

### Probing the Effects of Phasic LC Activation on Taste-Evoked GC Activity

To determine how phasic LC activation influences taste-evoked responses in GC, we expressed GCaMP8m (CaMKIIα-driven) in GC excitatory neurons and the excitatory opsin ChrimsonR (Flp-dependent) selectively in NE-producing LC neurons of *Dbh*^Flpo^ mice^17^ (Figure 1A). A relay lens was implanted over GC for miniscope calcium imaging, and an optical fiber was positioned above the ipsilateral LC for optogenetic stimulation. ChrimsonR-expressing LC axons densely innervated GC and were intermingled with GCaMP8m-expressing neurons (Figure 1B–C).

**Figure 1.**
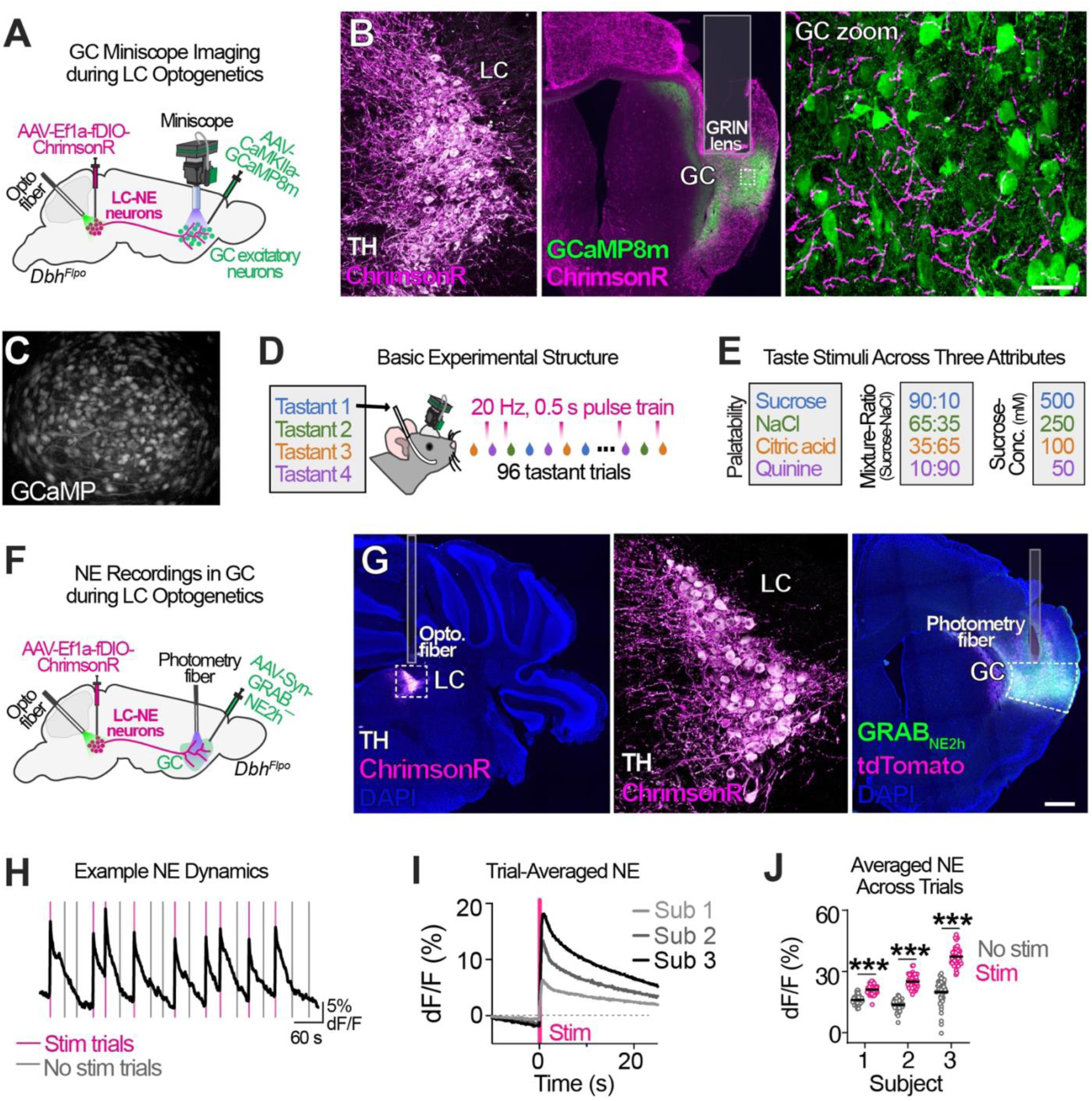
Approaches for Assessing Phasic LC Modulation of Taste-Evoked GC Responses. (A) Schematic of the viral-genetic approach and GC calcium imaging during LC optogenetic stimulation. (B) *Left*: LC neurons expressing the excitatory opsin ChrimsonR-tdTomato. *Middle*: ChrimsonR-positive LC axons project broadly, including to GC (dashed outline), where the GRIN lens is positioned (grey outline). *Right:* Zoomed view of GC showing excitatory neurons expressing GCaMP8m. Scale bars: 101.7 µm (left), 789 µm (middle), 64.4 (right). (C) Example calcium-imaging field of view. (D) *Experimental design*. Mice received one of four pseudo-randomly delivered taste stimuli via intraoral cannulae; LC neurons were phasically stimulated (20 Hz, 0.5s pulse train) before taste delivery on half the trials. (E) Taste stimuli used to assess three taste attributes. (F) Schematic of the viral-genetic strategy and NE fiber photometry during LC optogenetic stimulation. (G) *Left*: Coronal hindbrain section showing ChrimsonR-tdTomato expression in LC neurons; dashed line outlines the LC and the grey line marks the optical-fiber placement. *Middle*: Higher-magnification view of ChrimsonR-expressing LC neurons. *Right*: Coronal forebrain section showing GC expression of GRAB_NE2h_ and tdTomato (used for motion-artifact correction); dashed line outlines GC and the grey line marks optical-fiber placement. Scale bars: 498 µm (left), 66.7 µm (middle), 500 µm (right). (H) Example fiber-photometry trace showing NE dynamics induced by phasic LC stimulation (20 Hz, 0.5 s). Vertical lines denote the example timing of tastant delivery; note that no tastant was delivered in this validation experiment. (I) Trial-averaged NE fluorescence aligned to stimulation onset. *n*=3 mice. (J) NE fluorescence averaged over 1-s epochs for phasic-stimulation and no-stimulation conditions across subjects. Two-way ANOVA: Stimulation, *F*_2,282_*=*648.6, *P*<0.0001; Stimulation x subjects, *F*_2,282_*=*63.19, *P*<0.0001; Subjects, *F*_2,282_*=*210.7, *P*<0.0001. Šídák’s post-hoc test: *\*\*\*P*<0.001, Stim vs. No stim. Dots show mean responses for *n=*48 trials/condition (3 mice); horizontal lines show the mean per condition.

To precisely control stimulus delivery, mice were implanted with an intraoral cannula (IOC) (Figure 1D). Tastants were delivered through four independent microtubes inserted into the IOC, preventing cross-stimulus mixing. Each session consisted of 96 trials with repeated presentations of four stimuli (Figure 1D). On half of the trials, tastant delivery was preceded by a brief phasic LC stimulation (0.5 s; 20 Hz, 5-ms pulses), ending 0.5 s before taste delivery. This stimulation timing was selected to mimic phasic LC activation that immediately precedes a feeding bout^45^, which may prime the processing of taste information. All recordings were performed in awake, water-restricted mice. To test whether LC-dependent modulation varied across sensory dimensions, we tested three stimulus sets that differed along defined attributes: palatability (sucrose > NaCl > citric acid > quinine), mixture composition (four sucrose–NaCl ratios), and intensity (four sucrose concentrations) (Figure 1E).

To verify that our stimulation protocol effectively increased LC output to GC, we performed validation experiments in separate *Dbh*^Flpo^ mice expressing the NE sensor GRAB_NE2h_^46^ in GC and Flp-dependent ChrimsonR in the ipsilateral LC (Figure 1F–G). Using spectrally resolved fiber photometry^45,47,48^, we measured NE dynamics in GC using the same stimulation timing as in miniscope experiments but without tastant delivery (Figure 1D). Brief phasic LC stimulation reliably evoked robust increases in NE release across mice (Figure 1H–I). Although NE fluorescence did not fully return to baseline within the 25–35 s inter-trial interval, this prolonged decay was likely exaggerated by the off-kinetics of GRAB_NE2h_ (𝜏_off_ = 1.93 s^46^), as NE transients evoked by brief LC or NE pathway stimulation typically return to baseline within 10 seconds when measured using fast-scan cyclic voltammetry^49–53^. Thus, the prolonged GRAB_NE2h_ fluorescence decay likely reflects sensor kinetics rather than sustained NE elevation throughout the inter-trial interval. Notably, NE fluorescence signals during stimulation (stim) trials were clearly distinguishable from the no-stimulation (no-stim) trials (Figure 1J). These results confirm the efficacy of our stimulation protocol and justify comparisons of GC activity between these conditions.

### Phasic LC Activation Bidirectionally Modulates Taste-Evoked GC Responses

We first examined the effects of phasic LC activation on GC responses to four basic taste stimuli that vary in palatability. We identified neurons exhibiting tastant-dependent responses in stim and/or no-stim trials (hereafter termed *taste-discriminative* neurons). Among these, 20.8% showed differential responses in stim and no-stim trials for at least one tastant; these were classified as *LC-modulated* neurons (Figure 2A). In control mice expressing tdTomato in LC, only 2.88% of taste-discriminative neurons met criteria for LC modulation (Figure S1), suggesting that the observed response differences primarily reflect LC activation rather than photostimulation artifacts or false positives.

**Figure 2.**
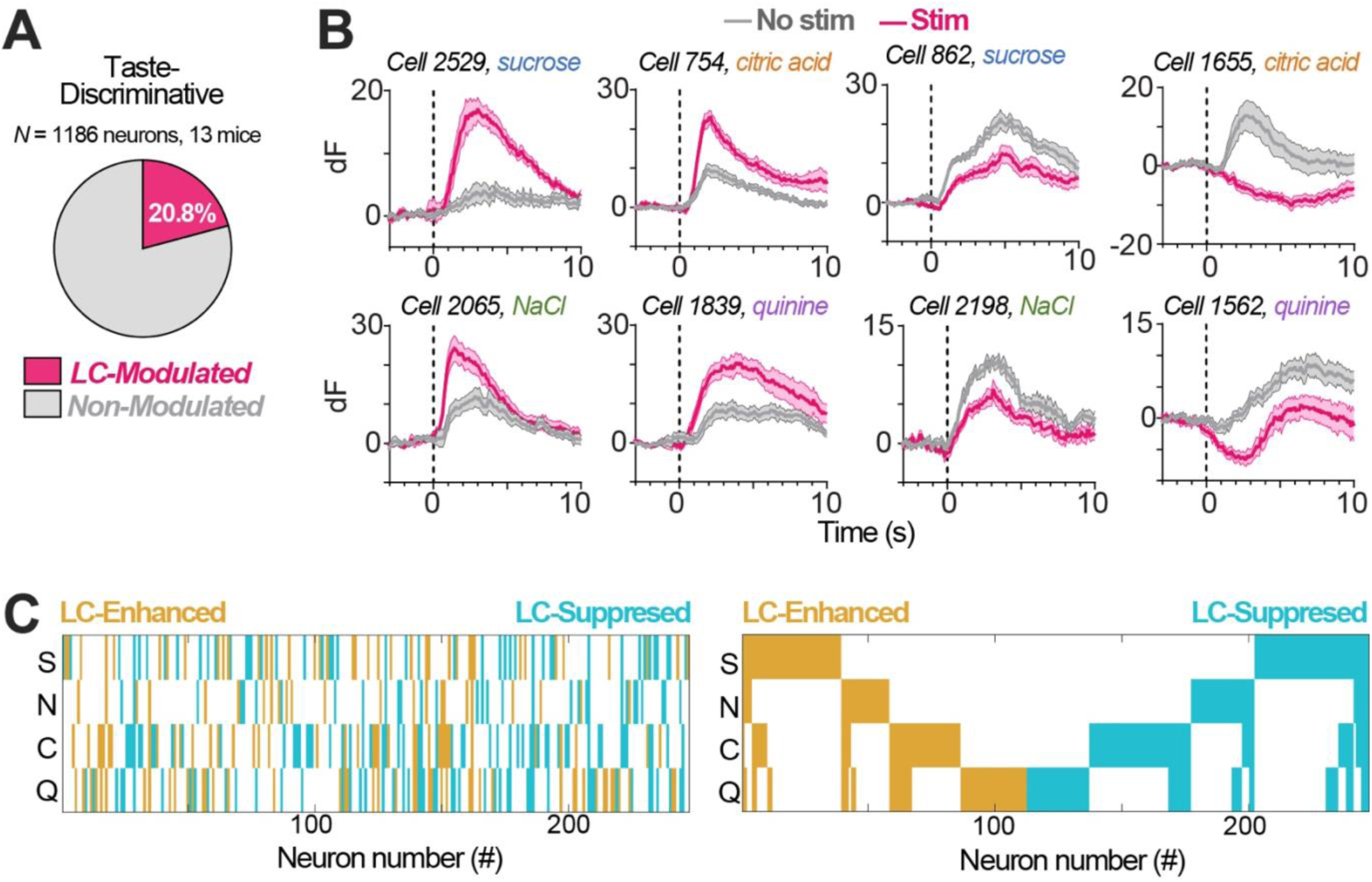
Phasic LC Activation Bidirectionally Alters Taste-Evoked Responses. (A) Proportion of taste-discriminative GC neurons showing significant modulation by LC stimulation. (B) Average responses from eight representative GC neurons, each responding to a single tastant (sucrose, NaCl, citric acid, or quinine), in the presence and absence of LC stimulation (taste delivery at *t*=0 s). Shading indicates ±SEM. (C) *Left*: Modulation direction for all LC-modulated GC neurons. Each tick shows the effect of LC activation on a neuron’s response to sucrose (S), NaCl (N), citric acid (C), or quinine (Q). Colors indicate LC-enhanced (gold) or LC-suppressed (teal) cell-taste pairs. *Right*: The same plot, sorted by modulation direction.

Consistent with findings in other LC target regions^18–20^, phasic LC activation produced both enhancement and suppression of GC taste responses (Figure 2B). In some cases, LC modulation gated GC activity by recruiting previously non-responsive neurons (*cell 2529*), or by suppressing excitatory responses, occasionally resulting in sign reversals (*cell 1655*) (Figure 2B). Across all significant cell–tastant pairs, individual neurons could be modulated for multiple tastants; however, the direction of modulation within a given neuron was uniformly enhancing or suppressive (Figure 2C). The proportions of LC-enhanced and LC-suppressed neurons were approximately balanced.

### Phasic LC Activation Enhances GC Palatability Correlation Without Altering Tastant Separability

To determine how LC activation alters GC taste encoding, we analyzed a pseudo-population of LC-modulated, taste-discriminative neurons pooled across mice. To assess population-level taste discriminability, we trained linear decoders to classify tastants using population dynamics. Decoding accuracy increased monotonically with population size; however, accuracy did not differ significantly between stim and no-stim trials (Figure 3A), suggesting that phasic LC activation does not alter overall tastant discriminability in GC.

**Figure 3.**
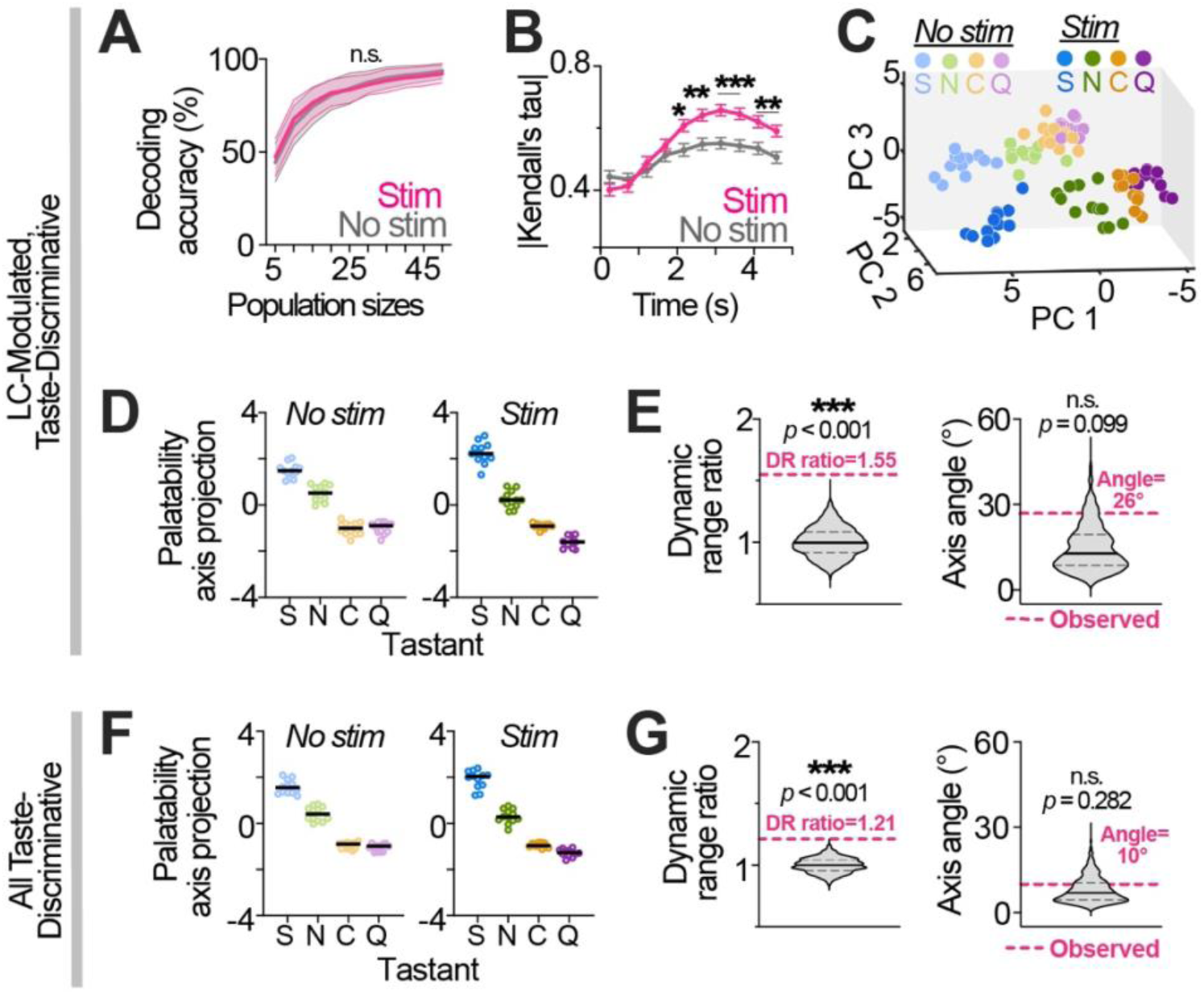
Phasic LC Activation Impacts GC Representation of Taste Palatability. (A) Decoding accuracy for LC-modulated, taste-discriminative GC neurons across population sizes. Data are mean ± SD of bootstrapped means. Permutation tests: n.s. at all population sizes. n.s., non-significant. (B) Correlations between response magnitude and palatability rank across LC-modulated neurons and time bins. Two-way RM ANOVA (time × stimulation): *F*_5.766, 1418_= 6.259, *P*<0.0001. Main effects—stimulation: *F*_1.000, 246.0_= 13.41, *P*=0.0001; time: *F*_5.112, 1258_= 33.98, *P*<0.0001. Bonferroni post hoc: \*\*\**P*< 0.001, \*\**P*<0.01, \**P*<0.05. Values are mean ± SEM (247 cells). (C) Population responses of LC-modulated GC neurons to tastants projected into principal-component space with and without stimulation. *n=*12 trials/condition. Sucrose (S); NaCl (N); citric acid (C); quinine (Q). (D) Population responses of LC-modulated neurons projected onto the respective palatability axis for stimulation (right) and no stimulation (left) trials. *n=*12 trials/condition. (E) Permutation tests for dynamic-range (DR) ratio (left) and palatability-axis angle (right) for LC-modulated neurons. DR is the Euclidian distance between the mean responses to S and Q. The DR ratio compares stimulation to no stimulation. Violin plots show the distributions of permuted values (null distribution), with median (black solid lines) and quartiles (black dashed). Magenta lines mark observed values. (F) Population responses of all taste-discriminative neurons projected onto the respective palatability axis for stimulation (right) and no stimulation (left). *n=*12 trials/condition. (G) Permutation tests for DR ratio (left) and palatability-axis angle (right) for all-taste discriminative neurons.

We next examined palatability encoding using a standard four-tastant panel^39,54–57^. These tastants differ in hedonic value to mice, with the expected ranking of sucrose > NaCl > citric acid > quinine, which we confirmed using a brief-access taste test (Figure S2). To quantify palatability-related information at the single-neuron level, we calculated the absolute rank correlation (Kendall’s tau) between each neuron’s mean response to the four tastants and their palatability ranking^37,58^ (Figure 3B). Palatability correlation increased gradually following tastant delivery and peaked around 3 s. This time course is delayed relative to prior electrophysiology studies^36,39^, in which palatability coding peaks ∼1–2.5 s after taste delivery, likely reflecting the slower kinetics of calcium imaging. Among LC-modulated neurons, mean palatability correlation was significantly higher in stim than no-stim trials, demonstrating that phasic LC activation enhances palatability coding at the single-neuron level.

### Phasic LC Activation Stretches GC Dynamic Range Along a Palatability-Relevant Axis

To examine how LC activation reshapes palatability coding at the population level, we performed principal component analysis (PCA) on evoked responses of LC-modulated neurons. Tastant responses formed a low-dimensional manifold organized primarily along a palatability-related axis (Figure 3C). We approximated the palatability axis separately for stim and no-stim conditions as vectors connecting the centroids of palatable (sucrose, NaCl) and aversive (citric acid, quinine) tastants. The most prominent effect of phasic LC activation was a coherent shift in the response manifold that was largely orthogonal to the palatability axes (Figure 3C), which may reflect changes in other GC coding directions (see *Discussion*).

To isolate LC-induced changes related to palatability coding, we projected responses onto their respective palatability axes (Figure 3D). Phasic LC activation increased the separation between sucrose and quinine—i.e., the dynamic range of the palatability axis—by 1.55-fold. A permutation test confirmed that this increase significantly exceeded that expected from trial-to-trial variability (Figure 3E). We next conducted permutation tests on the angle between stim- and no-stim-derived palatability axes and found no change in axis orientation (Figure 3E). Both the dynamic range expansion and axis stability were also observed when analyses were extended from LC-modulated neurons to the broader population of taste-discriminative neurons (Figure 3F–G).

Examination of pairwise tastant distances revealed that this expansion was non-uniform and was driven primarily by increased separation between sucrose and the remaining tastants (Figure S3A). Because the palatability axis orientation remained stable across conditions, we projected responses from both conditions onto a common (no-stim-derived) palatability axis to determine tastant-specific shifts. We found that phasic LC modulation shifted NaCl, citric acid, and quinine responses toward a more aversive direction, while leaving sucrose responses largely unchanged (Figure S4), thereby accounting for the non-uniform expansion of palatability dynamic range.

To determine whether LC modulation broadly expanded the stimulus manifold beyond the palatability dimension, we quantified the mean perpendicular distance of responses from the palatability axis, referred to as dispersion. Phasic LC activation did not significantly alter dispersion relative to permuted values, though it showed a trend toward expansion (*p* = 0.074; Figure S5A).

Together with the non-uniform changes in pairwise tastant distances, these findings indicate that LC modulation does not simply scale the population representation isotropically. Procrustes analysis further showed that LC modulation reorganized tastant centroids into a configuration more closely aligned with palatability ordering (Figure S5B–C).

### Both LC-Enhanced and LC-Suppressed Neurons Contribute to Enhanced Palatability Coding

Given that LC-modulated neurons can be subdivided into LC-enhanced and LC-suppressed subtypes, we next examined how each subtype contributes to changes in GC taste encoding. Although neither subtype showed changes in overall decoding accuracy, both exhibited significant increases in palatability correlation under phasic LC activation (Figure S6A, D).

Further, both subtypes had significant increases in dynamic range along the palatability axis (Figure S6B–C, E–F), demonstrating their role in palatability-axis elongation. Notably, axis rotation was observed in the LC-enhanced (Figure S6C) but not in LC-suppressed neurons (Figure S6F).

### Phasic LC Activation Stretches and Rotates GC Representations of Mixture Ratios

Given the role of GC in encoding mixture composition^59,60^, we next examined how phasic LC activation affects GC encoding of mixtures of sucrose–NaCl (90:10, 65:35, 35:65, 10:90). Because sucrose and NaCl differ in palatability (Figure S2), these mixtures likely vary in both mixture-ratio and palatability. Consistent with the palatability experiment, 20.6% of mixture-ratio-discriminative neurons were LC-modulated (Figure 4A), including both LC-enhanced and LC-suppressed subtypes (Figure 4B). Focusing on the LC-modulated neurons, we found that mixture-ratio decoding accuracy did not significantly differ between stim and no-stim trials (Figure 4C), indicating that phasic LC activation did not alter overall mixture separability. In contrast, LC activation significantly increased the correlation between neuronal responses and mixture–ratio ranking (Figure 4D), suggesting that LC modulation enhances GC encoding of graded differences across the sucrose–NaCl mixtures.

**Figure 4.**
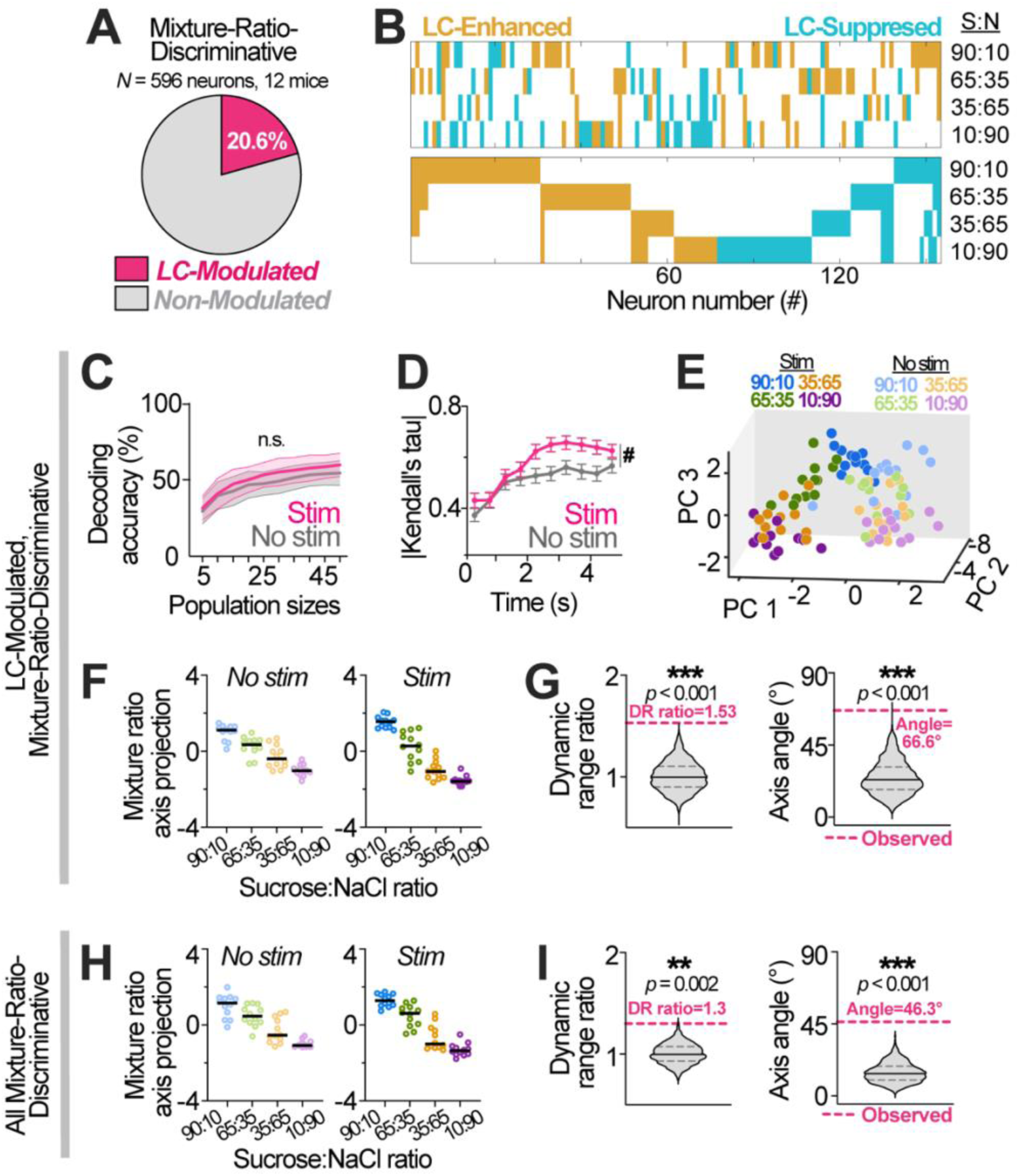
Phasic LC Activation Impacts GC Representation of Mixture Ratio Composition. (A) Proportion of mixture-discriminative GC neurons significantly modulated by LC stimulation. (B) *Top*: Modulation direction for LC-modulated GC neurons. Each tick shows the effect of LC activation on a neuron’s response to different sucrose (S)–NaCl (N) mixture ratios, with colors indicating modulation direction. *Bottom*: The same data, sorted by modulation direction. |(C) Decoding accuracy for LC-modulated, mixture-ratio-discriminative GC neurons across population sizes. Data are mean ± SD of bootstrapped means. Permutation tests: no significant effects at any population size. (D) Correlations between response magnitude and mixture-ratio rank across LC-modulated neurons and time bins. Two-way RM ANOVA: Stimulation: *F*_1.000, 122.0_= 6.169, *P*=0.0144; time: *F*_5.565, 679_= 20.79, *P*<0.0001. Time × stimulation: *F*_5.128, 625.7_= 1.597, *P*=0.1570. ^#^*P*<0.05, main effect of stimulation. Values are mean ± SEM (123 cells). (E) Population responses of LC-modulated GC neurons to different sucrose:NaCl ratios, projected into PCA space with and without stimulation. *n=*12 trials/condition. (F) Population responses of LC-modulated neurons projected onto the respective mixture-ratio axis for stimulation (right) and no stimulation (left) trials. *n=*12 trials/condition. G) Permutation tests for dynamic-range (DR) ratio (left) and mixture-ratio-axis angle (right) in LC-modulated neurons. DR is the Euclidian distance between mean responses to 90:10 and 10:90 sucrose:NaCl mixtures. The DR ratio compares stimulation vs. no stimulation. Violin plots show the distributions of permuted values (median: solid black; quartiles dashed); magenta lines indicate observed values. (H) Population responses of all mixture-ratio-discriminative neurons projected onto the respective mixture-ratio axis for stimulation (right) and no stimulation (left). *n=*12 trials/condition. (I) Permutation tests for DR ratio (left) and mixture-ratio-axis angle (right) for all mixture-ratio-discriminative neurons.

To characterize population-level transformations, we applied PCA to GC mixture-ratio responses (Figure 4E). We defined a mixture-ratio axis by connecting the 90:10 and 10:90 centroids and projected responses onto the respective axis. Phasic LC activation increased the dynamic range along this axis (Figure 4F–G). Pairwise mixture-distance analysis revealed that this expansion was non-uniform, primarily driven by increased separation between mixtures with larger differences in composition (Figure S3B). LC activation also induced a rotation of the mixture axis (Figure 4G). This rotation may reflect unequal stretching of latent dimensions encoding *pure* mixture composition and palatability, though alternative explanations remain possible (see *Discussion*). Both stretching and axis rotation persisted when analyses included all mixture-ratio-discriminative neurons (Figure 4H–I), showing that LC-induced geometric reorganization is robust when considering the broader population.

Dispersion analysis showed a trend toward reduced representational distance in dimensions orthogonal to the mixture axis (Figure S5D). Together with increased dynamic range, this pattern supports selective elongation of the stimulus manifold along a mixture-relevant axis, a transformation visualized using Procrustes analysis (Figure S5E–F).

To further dissect subtype contributions, we analyzed LC-enhanced and LC-suppressed neurons separately. LC-enhanced neurons exhibited both a non-significant trend toward improved decoding and significant increase in mixture-ratio rank correlation (Figure S7A). Conversely, LC-suppressed neurons showed no such effects (Figure S7D). PCA revealed dynamic range stretching and axis rotation in both subtypes (Figure S7B–C, E–F). Thus, LC-induced transformations of mixture representations were attributable to both modulatory subtypes.

### Phasic LC Activation Modestly Expands and Rotates GC Representations of Sucrose Concentration

We next examined LC’s effect on GC encoding of tastant intensity using four sucrose concentrations (50, 100, 250, and 500 mM). Because sucrose palatability increases monotonically with concentration^61,62^, these stimuli likely vary in both intensity and palatability, analogous to mixtures. Consistent with prior results, 19.3% of sucrose-concentration-discriminative neurons were LC-modulated, comprising both LC-enhanced and LC-suppressed subtypes (Figure 5A–B).

**Figure 5.**
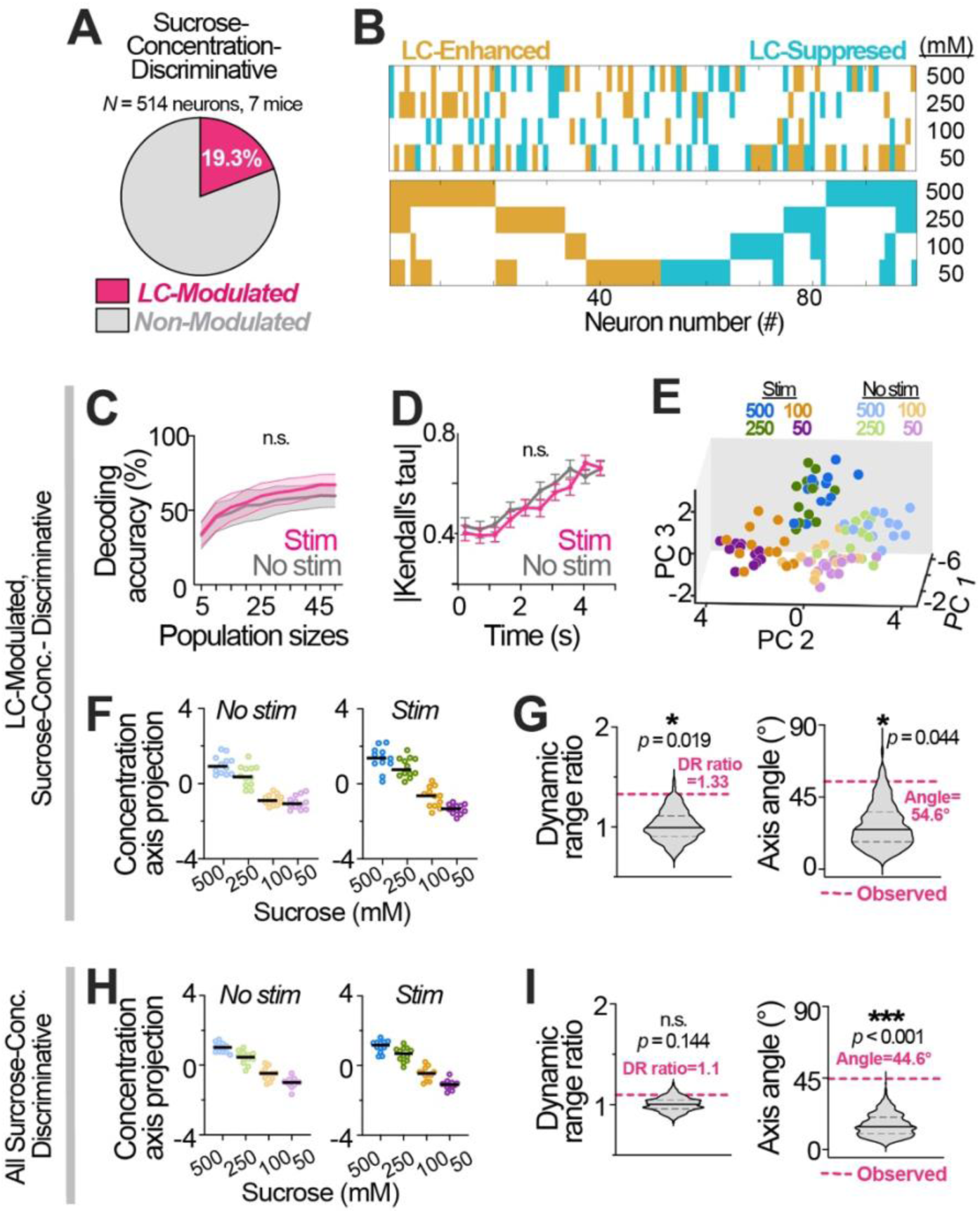
Phasic LC Activation Impacts GC Representation of Sucrose Intensity. (A) Proportion of sucrose-concentration-discriminative GC neurons significantly modulated by LC stimulation. (B) *Top*: Modulation direction for all LC-modulated GC neurons. Each tick shows the effect of LC activation on a neuron’s response to different sucrose concentrations (mM), with colors indicating modulation direction. *Bottom*: The same data, sorted by modulation direction. (C) Decoding accuracy for LC-modulated, sucrose-concentration-discriminative GC neurons across population sizes. Data are mean ± SD of bootstrapped means. Permutation tests show no significant effects at any population size. (D) Correlations between response magnitude and sucrose-concentration rank across LC-modulated neurons and time bins. Two-way RM ANOVA: Stimulation: *F*_1.000, 98.0_= 0.7657, *P*=0.3837; Time: *F*_4.870, 477.2_= 26.38, *P*<0.0001. Time × Stimulation: *F*_5.594, 548.3_= 1.111, *P*=0.3543. No significant main effect of stimulation. Values are mean ± SEM (99 cells). (E) Population responses of LC-modulated GC neurons to different sucrose concentrations, projected into principal-component space with and without stimulation. *n=*12 trials/condition. (F) Population responses of LC-modulated neurons projected onto the respective sucrose-concentration axis for stimulation (right) and no stimulation (left) trials. *n=*12 trials/condition. (G) Permutation tests for the dynamic-range (DR) ratio (left) and sucrose-concentration-axis angle (right) in LC-modulated neurons. DR is the Euclidian distance between mean responses to 500 mM and 50 mM sucrose. The DR ratio compares stimulation vs. no stimulation. Violin plots show the permuted (null) distributions (median: solid black; quartiles dashed). Magenta lines mark observed values. (H) Population responses of all sucrose-concentration-discriminative neurons projected onto the respective concentration axis for stimulation (right) and no stimulation (left) trials. *n=*12 trials/condition. (I) Permutation tests for DR ratio (left) and sucrose-concentration-axis angle (right) for all concentration-discriminative neurons. Violin plots show the distributions of permuted values (median: solid black; quartiles dashed); magenta lines for observed values.

Among LC-modulated neurons, phasic LC activation did not significantly alter decoding accuracy of sucrose concentration (Figure 5C) or correlation between neuronal responses and concentration (Figure 5D), indicating limited modulation of intensity coding. Nevertheless, PCA revealed a stimulation-dependent increase in dynamic

range along the sucrose-concentration axis (Figure 5E–G). Pairwise concentration-distance analysis revealed modest changes, with only the 250 versus 50 mM comparison surviving multiple-comparison correction (Figure S3), consistent with relatively weak dynamic-range expansion. LC modulation also induced significant rotation of the concentration axis (Figure 5G), possibly reflecting differential stretching of latent axes encoding pure concentration and palatability. When analyses were extended to all sucrose-concentration-discriminative neurons, axis rotation persisted, whereas dynamic-range stretching was no longer significant (Figure 5H–I), suggesting that axis rotation is the more robust population-level effect of LC modulation. Dispersion analysis revealed significant LC-induced expansion in dimensions orthogonal to the concentration axis (Figure S5G–I), possibly due to non-linearity in concentration coding not captured by the approximated concentration axis.

We next examined the contributions of LC-enhanced and LC-suppressed subtypes and found no effects in either group on decoding accuracy or rank correlation under phasic LC activation (Figure S8A, D). In contrast, PCA revealed increased dynamic range and a trend toward axis rotation in LC-enhanced neurons (Figure S8B–C), with no effects in LC-suppressed neurons (Figure S8E–F), suggesting that LC-enhanced neurons are the primary contributors to LC-induced changes in sucrose representation.

### Parsing Tuning Modulations Underlying Population-Level Effects of Phasic LC Activation

The preceding analyses showed that phasic LC activation induces distinct geometric transformations of GC population activity, including (but not limited to) dynamic-range expansion and axis rotation (Figure 6A). Because representational geometry is determined by neuronal tuning^34^, we next examined the LC-induced tuning changes underlying these transformations. We defined each neuron’s tuning profile as its mean response across a tastant set and, using a model comparison approach, classified LC-induced changes as *scale*–*shift* modulation, *flexible* modulation, or no modulation for each experiment (Figure 6B). Scale–shift modulation reflects a uniform transformation of the tuning profile, captured by a common multiplicative factor and additive offset applied across all tastants. In contrast, flexible modulation reflects tastant-specific changes that cannot be explained by uniform scaling or shifting. Example scale–shift and flexible neurons from the palatability experiment are shown in Figure 6C and Figure 6D, respectively. Scale–shift modulation preserved the relative response pattern across tastants, whereas flexible modulation could reorganize this pattern, including changes in rank order. Across experiments, basic-taste-discriminative neurons showed more scale–shift modulation and less flexible modulation than mixture-ratio- and sucrose-concentration-discriminative neurons (Figure 6E, H, K; Figure S9A).

**Figure 6.**
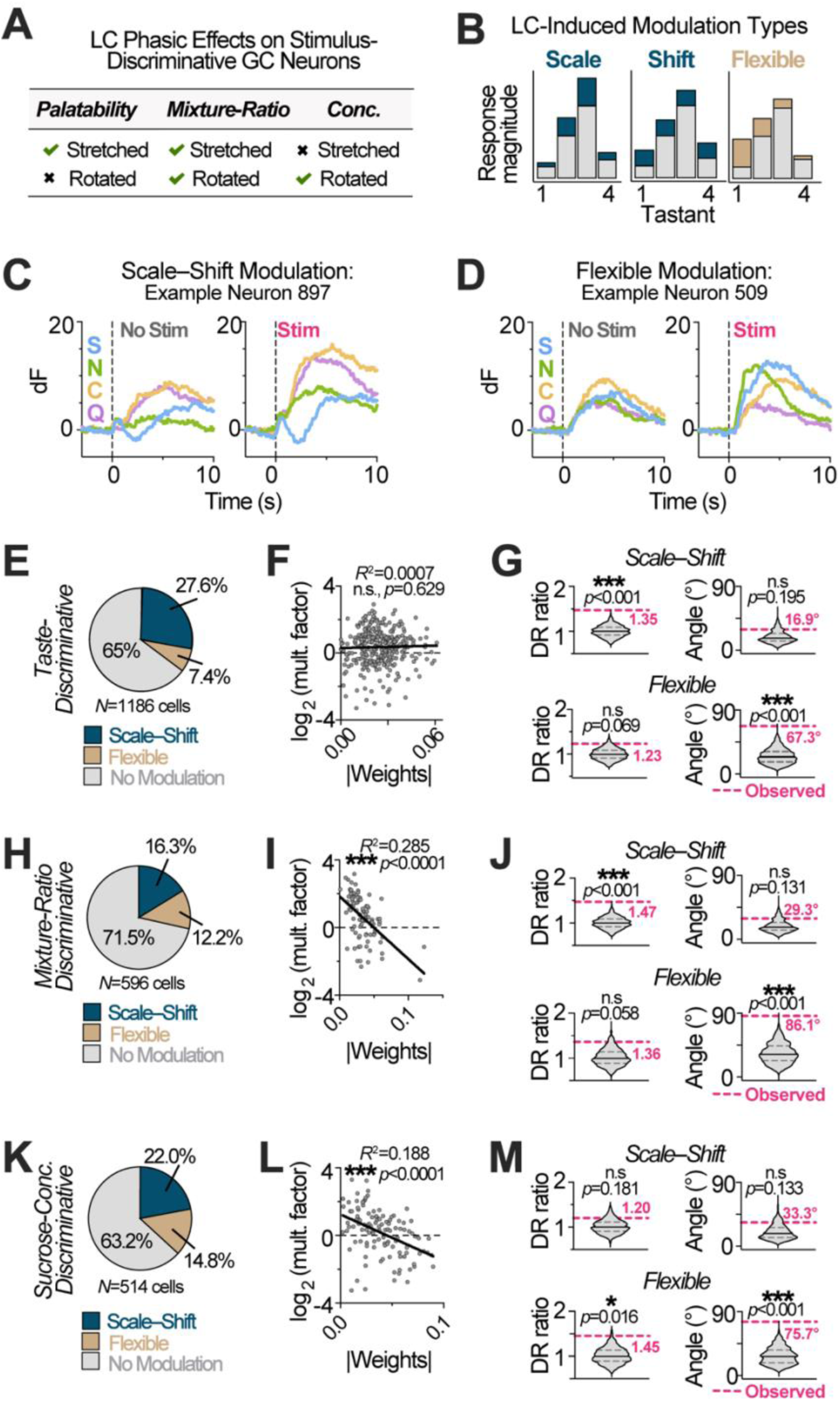
Phasic LC Activation Reshapes Neuronal Tuning in an Attribute-Dependent Manner. (A) Summary of LC-induced transformations on stimulus-discriminative GC neurons across attributes. (B) Schematic illustrating different types of modulation on neuronal tuning profile, including scaling (blue), shifting (blue), and flexible (gold) modulation. Grey shows stimulus tuning in the absence of modulation. (C-D) Example GC neurons showing scale–shift (C) or flexible modulation (D) during stimulation (right) and without stimulation (left) in the palatability experiment. Data are mean responses to each tastant: S, sucrose; N, NaCl; C, citric acid; Q, quinine. (E, H, K) Proportion of modulation types in GC neurons discriminative for palatability (E), mixture ratio (H), or sucrose concentration (K). (F, I, L) Linear regressions of absolute weights versus log multiplicative factor for scale–shift modulated GC neurons discriminative for palatability (F), mixture ratio (I), or sucrose concentration (L). Statistical results: Panel F: slope = 2.377 ± 4.909; *R*^2^ = 0.0007208; *F*_1, 325_= 0.2344, *P*=0.6286. Panel I: slope = -37.11 ± 6.038; *R*^2^ = 0.2845; *F*_1, 95_= 37.77, *P*<0.0001. Panel L: slope = -27.52 ± 5.421; *R*^2^ = 0.1884; *F*_1, 111_= 25.77, *P*<0.0001. (G, J, M) Permutation tests for the dynamic-range (DR) ratios (left) and axis angle (right) in scale–shift (top) and flexible (bottom) modulation types for palatability (G), mixture ratio (J), and sucrose concentration (M) experiments. Violin plots show permuted null distributions (median: solid black; quartiles dashed), with observed values in magenta.

### LC-Mediated Gain Modulation is Structured and Attribute-Specific

We next examined the additive and multiplicative factors associated with scale–shift-modulated neurons, given their potential contribution to population-level transformations. Additive factors shift the population manifold without changing its shape or size, whereas multiplicative factors can expand, compress, and/or rotate it. Both additive and log-transformed multiplicative factors spanned positive and negative values (Figure S9B). Median log-multiplicative factors were positive across all experiments, indicating an overall bias toward expansion (Figure S9C). Multiplicative effects were less expansive in the sucrose-concentration experiment than in the mixture experiment, consistent with the more modest expansion of sucrose dynamic range.

Given the relevance of multiplicative effects (gain modulation) to manifold expansion, we next examined whether multiplicative factors were related to each neuron’s contribution to the coding axis, measured by its absolute axis weight. No correlation was observed in the palatability experiment (Figure 6F), whereas significant negative correlations emerged in the mixture-ratio and sucrose-concentration experiments (Figure 6I, L). Thus, under phasic LC modulation, neurons contributing less strongly to these coding axes were preferentially expanded, whereas stronger contributors were relatively compressed.

### Distinct Contributions of Scale–Shift and Flexible Modulation to LC-Induced Transformations

We next applied PCA separately to scale–shift- and flexibly modulated neurons to determine their contributions to coding-space transformations. Scale–shift modulation increased dynamic range in the palatability and mixture-ratio experiments (Figure 6G, J), but not in the sucrose-concentration experiment (Figure 6M), consistent with the stronger expansion bias of multiplicative factors in the former two datasets (Figure S9C). Despite its potential to reweight neuronal contributions, scale–shift modulation did not induce axis rotation in any experiment. Thus, scale–shift modulation primarily contributed to LC-induced dynamic-range expansion.

In contrast, flexible modulation induced significant axis rotation across all experiments (Figure 6G, J, M), consistent with its ability to reshape neuronal tuning. Flexible modulation also increased dynamic range in the sucrose-concentration experiment, but not in the palatability or mixture-ratio experiments. Analysis of rank correlations further revealed that scale–shift modulation increased response–stimulus correlation in the palatability and mixture experiments (Figure S9D, F, H), whereas flexible modulation increased these correlations across all attributes (Figure S9E, G, I). Together, these findings indicate that scale–shift and flexible modulation make distinct contributions to LC-induced population transformations, with the former primarily supporting dynamic-range expansion and the latter driving axis rotation.

### Tonic LC Activation Has Limited Impact on GC Taste Encoding

Previous work suggests that phasic and tonic LC activation can produce qualitatively different effects on downstream circuits^26,28,63^. To examine the impact of tonic LC activation on GC taste encoding, we conducted three additional experiments using tastants varying in palatability, mixture ratio, and sucrose concentration, paired with an alternative photostimulation protocol. Tastant trials were divided into eight blocks, half of which included 5-Hz tonic LC stimulation (Figure 7A). Validation experiments confirmed that this protocol elicited robust NE release that persisted throughout each stimulation block (Figure 7B-C).

**Figure 7.**
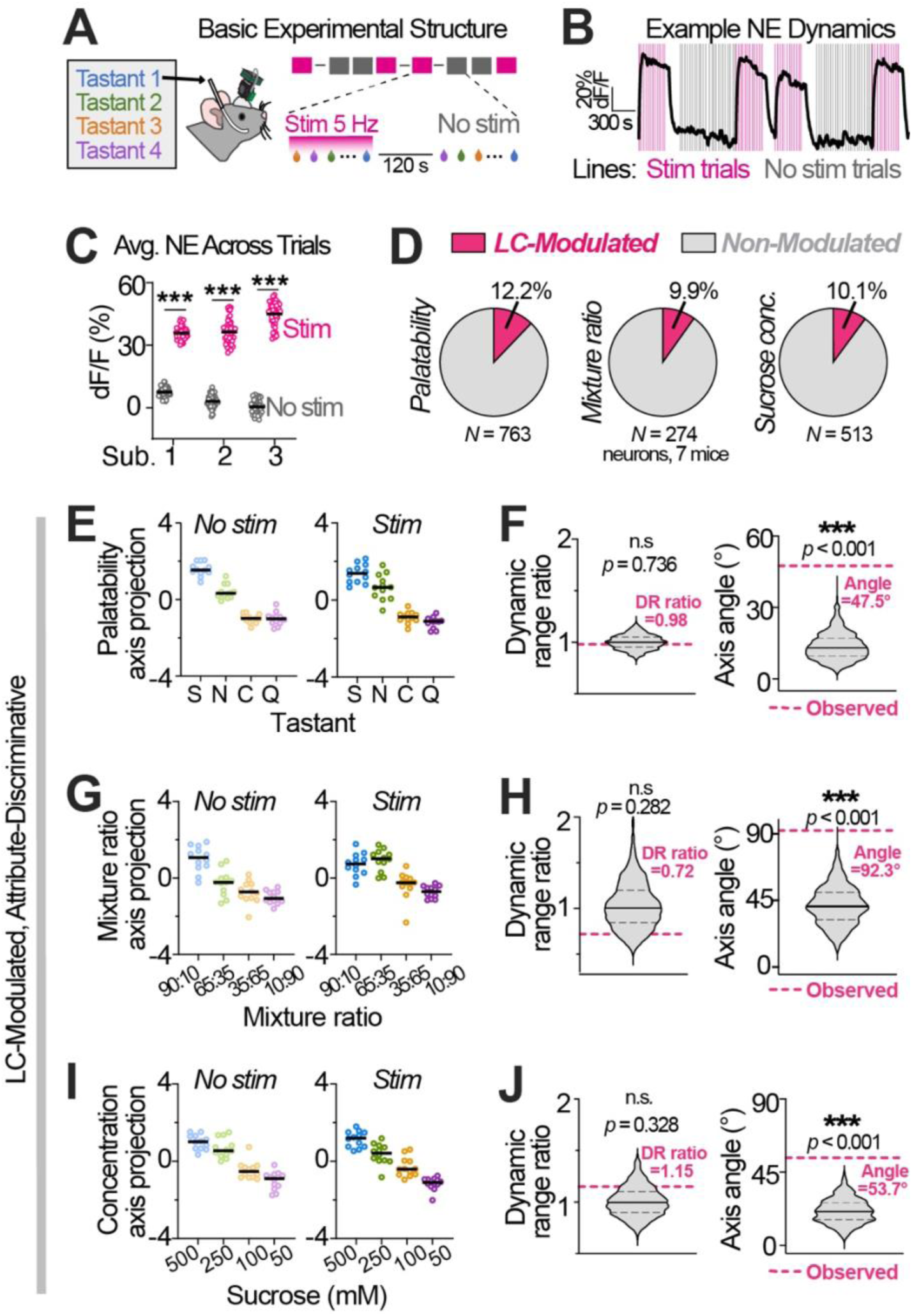
Effects of Prolonged Tonic LC Activation on GC Representation of Taste Attributes. (A) Mice received four pseudo-randomly delivered taste stimuli in eight blocks (12 deliveries per block). Tonic LC photostimulation (5 Hz) occurred in four blocks. The sequence of stimulation on (magenta) and off (grey) blocks are as depicted in the schematic. Each stimulation block is followed by a gap period (145–155 s) before the onset of the next block. (B) Example fiber-photometry trace showing NE dynamics evoked by tonic LC stimulation (5 Hz). Vertical lines denote the example timing of tastant delivery; note that no tastant was delivered in this validation experiment. (C) NE fluorescence averaged over 1-s epochs for tonic-stimulation and no-stimulation trials. Two-way ANOVA: Stimulation, *F*_2,282_*=*5392, *P*<0.0001; Stimulation x subjects, *F*_2,282_*=*97.60, *P*<0.0001; Subjects, *F*_2,282_*=*14.00, *P*<0.0001. Šídák’s post-hoc: *\*\*\*P*<0.001. Dots show *n=*48 trials/condition (3 mice); lines show means. (D) Proportion of stimulus-discriminative GC neurons modulated by tonic LC stimulation across taste attributes. (E, G, I) Population responses of LC-modulated neurons projected onto the palatability (E), mixture ratio (G) and sucrose concentration (I) axes for stimulation (right) and no-stimulation (left) trials (*n=*12 trials/condition). F, H, J) Permutation tests for dynamic-range (DR) ratios (left) and axis angles (right) for GC neurons whose responses were significantly modulated by tonic LC stimulation in the palatability (F), mixture ratio (H), and sucrose concentration (J) experiments. Violin plots show permuted distributions (median: solid black; quartiles dashed), with observed values in magenta.

Despite robust NE release, tonic LC activation modulated only ∼10–12% of stimulus-discriminative neurons across attributes (Figure 7D), approximately half the proportion observed under phasic stimulation, though still higher than in tdTomato controls (Figure S1B). Tonic activation did not affect palatability or mixture-ratio decoding but reduced decodability for sucrose concentration (Figure S10). Tonic activation also did not alter response–stimulus correlations for any taste attribute (Figure S10).

PCA showed that tonic LC activation produced axis rotation across all attributes without changing dynamic range (Figure 7E-J). When all stimulus-discriminative neurons were included, rotation persisted for mixture ratio and sucrose concentration, but not palatability (Figure S11). Together, these findings suggest that, unlike phasic activation, tonic LC activation produced modest transformations that are not clearly aligned with behaviorally relevant stimulus dimensions, suggesting a limited influence on GC taste representations.

## DISCUSSION

Here, we examined how LC activation shapes cortical taste encoding at both single-neuron and population levels. A central advance of this study is that it links LC activation patterns to specific geometric transformations of gustatory cortical population activity, revealing how a major neuromodulatory system can reorganize taste representations across multiple stimulus dimensions. Phasic LC activation (i) bidirectionally modulated taste-evoked responses, with each neuron showing a consistent direction of modulation, and (ii) enhanced correlations between neuronal responses and graded stimulus properties in the palatability and mixture-ratio experiments. At the population level, phasic LC activation (i) stretched the dynamic range of GC activity along attribute-relevant axes and (ii) rotated the mixture-ratio and sucrose-concentration axes, without affecting the palatability axis. Dynamic-range stretching was driven by structured multiplicative modulation, whereas axis rotation reflected flexible reshaping of neuronal tuning. In contrast to phasic LC activation, tonic activation influenced fewer GC neurons and produced axis rotation without dynamic-range stretching. These results indicate that brief LC activation reorganizes GC population coding to enhance stimulus representation, particularly along the behaviorally relevant palatability dimension.

### Population-Level Effects of LC Modulation

The geometry of population activity determines how sensory information is represented. Expansion along an attribute-relevant axis, for example, is linked to increased perceptual disparity of information along that dimension. Selective attention, for instance, can stretch neural representations along task-relevant axes^64,65^. Although LC activity is more commonly associated with global arousal than with top-down control, it may nonetheless bias information processing toward specific sensory attributes. Consistent with this idea, phasic LC activation primarily increased the separation between sucrose and other basic tastes along the palatability dimension without causing overall expansion in orthogonal dimensions, suggesting selective reorganization of palatability coding. The absence of expansion along other taste-differentiating dimensions may help explain the unchanged decoding performance under LC modulation. Our findings are consistent with prior work showing that behavioral state differentially influences taste identity and palatability representations in the GC^40,41^. Indeed, palatability differences may be more important for guiding feeding decisions than precise tastant identity. Notably, reorganization of palatability coding was driven primarily by aversive shifts in NaCl, citric acid, and quinine responses, whereas sucrose responses remained largely unchanged. The relative stability of sucrose representations may be related to its caloric value, a possibility that could be tested by comparing LC modulation of sucrose and non-caloric sweeteners. Together, these findings raise the possibility that LC modulation biases taste coding toward highly palatable or energy-rich foods (e.g., when navigating unfamiliar foraging environments), although this behavioral predication remains to be tested.

Axis rotation provides an additional transformation by which population geometry can be modulated. The rotation of mixture and sucrose-concentration axes admits several interpretations that do not require alterations in downstream readout. One possibility is that these axes reflect blending of two latent dimensions: objective stimulus properties and palatability. If LC modulation scales these dimensions unequally, the resulting attribute axis would rotate toward the dimension undergoing greater stretching. This account aligns with our observation that multiplicative modulation expands low-contribution neurons while compressing high-contribution neurons. If these groups preferentially represent distinct latent dimensions, such opposing gain modulation would reweight those dimensions. Alternatively, LC activation may recruit previously non-discriminative neurons into stimulus encoding, effectively pulling the attribute axis toward the subspace they span. These neurons would likely be classified as flexibly modulated, consistent with their role in both driving axis rotation and increasing response–stimulus correlations. Importantly, axis estimates could be sensitive to noise, particularly when the defining centroids are close together, and thus the precise angle should not be interpreted literally. This variability is addressed here by assessing significance through permutation testing.

Beyond stretching and rotation, phasic LC activation imposed a sizable, coherent shift of the response manifold largely orthogonal to the palatability axis, consistent with additive modulation of neuronal tuning. Although this shift could occur along another taste-related coding dimension, we did not observe large displacements aligned with mixture-ratio or sucrose-concentration axes. Alternatively, the LC-induced shift orthogonal to the palatability axis may reflect an internal-state dimension, such as arousal, that is linked to LC output^66,67^. This shift may also represent causal or compensatory changes in the network that enable or accompany expansion and/or rotation in the taste-relevant coding dimension.

### Mechanisms of LC-Induced Neuronal Effects

LC modulation can shape cortical taste responses through adrenergic receptor-mediated effects on (i) GC neurons, (ii) local recurrent GC circuits, and/or (iii) other brain regions influencing GC. Both glutamatergic and GABAergic GC neurons express transcripts for several adrenergic receptors, including the α1- and α2-receptor subtypes and β1-receptors^68^. Activation of these receptors can influence intrinsic and synaptic properties of cortical neurons^69–73^, potentially contributing to the enhancement, suppression, and diverse tuning effects observed here. LC output may also influence neuronal responses indirectly through its effects on other cortical microcircuit components, including inhibitory neurons, which were not examined in this study. Additionally, changes in GC activity may reflect LC’s impact on other nodes of the taste pathway, such as gustatory thalamus and basolateral amygdala, which primarily convey chemosensory^74–76^ and hedonic information^55,77,78^, respectively. Thus, our results may reflect both local adrenergic modulation within GC and LC-driven changes across the broader gustatory network. How LC influences GC neurophysiology and inter-regional interactions remains an important open question.

Characterizing whether LC enhances or suppresses neuronal responses to different tastants alone is insufficient to explain the observed reorganization of taste coding. Population-level effects such as palatability-axis expansion and coding-axis rotation depend on how tuning profiles are reshaped across stimuli. To examine LC-mediated tuning changes, we examined scale-shift and more flexible modulatory effects. Decomposing modulation into additive and multiplicative components (scale–shift) is a common framework for characterizing modulatory effects on neuronal tuning^79–81^. Multiplicative (gain) modulation, in particular, is a well-established mechanism by which the LC–NE system can facilitate cognition and information processing^11,82–84^. Although the cellular mechanisms underlying additive and multiplicative modulation have been extensively studied^81,85,86^, the basis of flexible modulation remains less clear, given its heterogeneous nature. Recruitment of coding neurons—a form of flexible modulation described above—may reflect unmasking of attribute-relevant inputs. Alternatively, flexible modulation may arise from differential scaling of narrowly tuned, labeled-line-like inputs, effectively altering the relative contribution of distinct sensory streams. Importantly, our classification of modulation types is descriptive and does not imply discrete mechanistic categories. Rather, this framework provides a way to relate diverse single-neuron tuning changes to the broader reorganization of GC population coding induced by LC activation.

### Links to State-Dependent Modulation

Prior work has revealed an enhancement in GC palatability coding during task disengagement^40^ and LiCl-induced sickness^41^. These behavioral states are accompanied by shifts in cortical state, reflected in local field potential patterns. Because LC–NE activity is tightly coupled to cortical states^87,88^ and can rapidly modulate it^89–92^, LC signaling provides a mechanistic link between cortical states, internal states, and sensory coding. Whereas these prior studies examined behavioral states lasting tens of minutes, our results showed that gustatory representations can be reorganized on the timescale of transient LC activation, contributing to moment-to-moment variability in sensory responses^4^.

Understanding LC function requires moving beyond broad labels such as “arousal” to consider precise LC dynamics in specific sensory-guided behaviors. In the context of taste, prior work shows that LC phasic activation occurs immediately before consumption^45^ and during brief gaps between lick bouts^48^. These dynamics—mimicked by our phasic stimulation pattern—suggest that LC activity may transiently prime GC before each influx of gustatory input. Notably, LC activity is suppressed *during* consumption bouts^12,45,48^, potentially stabilizing sensory representations by reducing LC-driven fluctuations. Moreover, many learning paradigms involve delivery of gustatory rewards immediately following reward-seeking actions. Thus, phasic LC activation accompanying these actions^13,14,93^ may extend into reward processing, thereby influencing learning.

The distinct effects of phasic and tonic LC activation observed here highlight the importance of activation pattern in neuromodulation. A classic framework of LC function proposes that moderately aroused states (associated with phasic LC activation) optimize information processing, whereas hyper-aroused states (associated with high tonic LC activation) do not^2,11,94^. While previous studies have shown that phasic and tonic LC activation can differentially affect sensory processing^26,28^, our results demonstrate that the sensory-enhancing effects of phasic activation in GC are absent under high tonic activation. Curiously, tonic LC activation reduced decodability of only sucrose-concentration-coding neurons, suggesting a selective diminishment in intensity representation. A proposed mechanism for divergent effects of LC activity patterns is that phasic activation preferentially engages high-affinity α1 receptors through moderate NE release, whereas sustained tonic activity recruits lower-affinity β receptors with distinct cellular consequences^95^. Consistent with our findings, recent fMRI work suggests that phasic LC activation produces brain-wide interaction patterns that may favor sensory processing relative to tonic activation^63^. Together, these findings suggest that LC’s influence on sensory coding depend on its activation pattern.

### Limitations and Open Questions

Several limitations should be considered when interpreting our findings. First, although our results suggest that phasic LC activation selectively reorganizes palatability representations, this interpretation is based on neural population geometry and will require behavioral validation. A definitive behavioral test will require an extensive, dedicated study combining optogenetic LC manipulation with palatability assays, including brief-access testing and quantitative taste-reactivity measurements, across appropriate controls, stimulation conditions, and sex-balanced cohorts, as these behaviors are known to exhibit sex differences^96–98^. Second, our interpretation of axis rotation as reflecting unequal stretching of covarying taste attributes (e.g., palatability vs. mixture-ratio or concentration) remains speculative. A more direct test would present the mixture-ratio or concentration stimulus sets together with the standard four-taste panel within the same session, allowing the latter to provide a stable palatability reference axis. Third, the specific effects observed in each experiment may depend on the tastant battery, including stimulus selection and concentration range. Finally, because our population-level analyses rely on distinct attribute axes defined in separate PCA spaces for each stimulus set, they do not readily permit formal statistical comparisons across experiments. Therefore, differences between palatability, mixture-ratio, and sucrose-concentration results should be interpreted cautiously.

Our work also provides a foundation for probing more advanced aspects of population coding, including response dynamics and noise correlations. In regard to dynamics, GC taste responses unfold as sequences of ensemble states that successively encode the presence, identity, and palatability of tastants^36,37,99^. How LC activity shapes these states and their transitions remains unknown and will likely require higher-temporal-resolution recordings such as electrophysiology. Another unexplored aspect is noise correlation—the shared trial-to-trial variability between neurons—which critically influences population information capacity^100–102^ and is regulated by internal states^103^ and neuromodulation^104^. Characterizing LC’s impact on noise correlations will require analyses of simultaneously recorded populations and larger numbers of trials than used here. Together, these open questions underscore the importance of the present study as a first causal framework for understanding how distinct temporal modes of LC activity reshape gustatory cortical representations. Rather than globally altering sensory responsiveness, phasic LC activation reorganizes the geometry of gustatory population activity, revealing a population-level mechanism through which LC shapes sensory representations. These findings also establish a foundation for future behavioral and circuit-level studies linking LC-mediated sensory coding to perception and behavior.

## STAR METHODS

Detailed methods are provided in the online version of this paper and include the following:

- KEY RESOURCES TABLE
- RESOURCE AVAILABILITY

`o` Lead Contact
`o` Materials Availability
`o` Data and Code Availability
- EXPERIMENTAL MODEL AND SUBJECT DETAILS
- METHOD DETAILS

`o` Survival Surgical Procedures
`o` Intracranial Viral Injections
`o` Intracranial Optical Fiber and GRIN Lens Implantation
`o` Intraoral Cannulae (IOC) Implantation
`o` Water Restriction Protocol
`o` Passive Tastant Delivery
`o` *In Vivo* Miniscope Calcium Imaging
`o` Optogenetic Stimulation Procedures
`o` Experimental Timeline
`o` Fiber Photometry Recordings of Photostimulation-Evoked NE Release
`o` Brief-Access Taste Testing
`o` Tissue Collection and Processing
`o` Immunohistochemistry
`o` Digital Image Processing
`o` Data Presentation and Visualization
- QUANTIFICATION AND STATISTICAL ANALYSIS

`o` Miniscope Movie Pre-Processing and Cell-Identification
`o` Identification of Taste-Discriminative and LC-Modulated Neurons
`o` Population Decoding
`o` Response–Stimulus Correlation
`o` Principal Component Analysis (PCA) and LC-Induced Transformations
`o` Modulation Type Analysis
`o` Fiber Photometry Data Analysis
`o` Brief-Access Taste Test Analysis

## SUPPLEMENTAL INFORMATION

Supplemental Figures S1-S10 can be found below.

## ACKNOWLEDGEMENTS

We thank Dr. Alfredo Fontanini for guidance on the intraoral delivery method. We also thank Leon Novak, Julia Chernyak, Anthony Esposito, and Abigail McFarland for assistance with genotyping, and Nikolas Supczak for histology support. We acknowledge the University of Connecticut’s Animal Care Services for animal husbandry support, the Advanced Light Microscopy Facility (RRID:SCR_027547) for imaging guidance, and the Mechanical/Glass Facility for custom intraoral connectors. This work was supported by the National Institutes of Health (R01 DC023564 and R00 DK119586), the Brain Research Foundation Seed Grant (BRFSG 2023-09), and UConn CLAS Research Equipment Fund and Startup Funds (to N.R.S.).

## AUTHOR CONTRIBUTIONS

Conceptualization: W.F., N.R.S.; investigation: W.F., P.T.; formal analysis: W.F.; data visualization: W.F., P.T., N.R.S.; writing – original draft: W.F., N.R.S.; writing - review & editing, W.F., P.T., N.R.S.; project administration, supervision and funding acquisition: N.R.S.

## DECLARATION OF GENERATIVE AI AND AI-ASSISTED TECHNOLOGIES IN THE MANUSCRIPT PREPARATION PROCESS

During the preparation of this work, the author(s) used ChatGPT for proofreading. The author(s) reviewed and edited the output as needed and take full responsibility for the content of the published article.

## STAR METHODS

### KEY RESOURCES TABLE

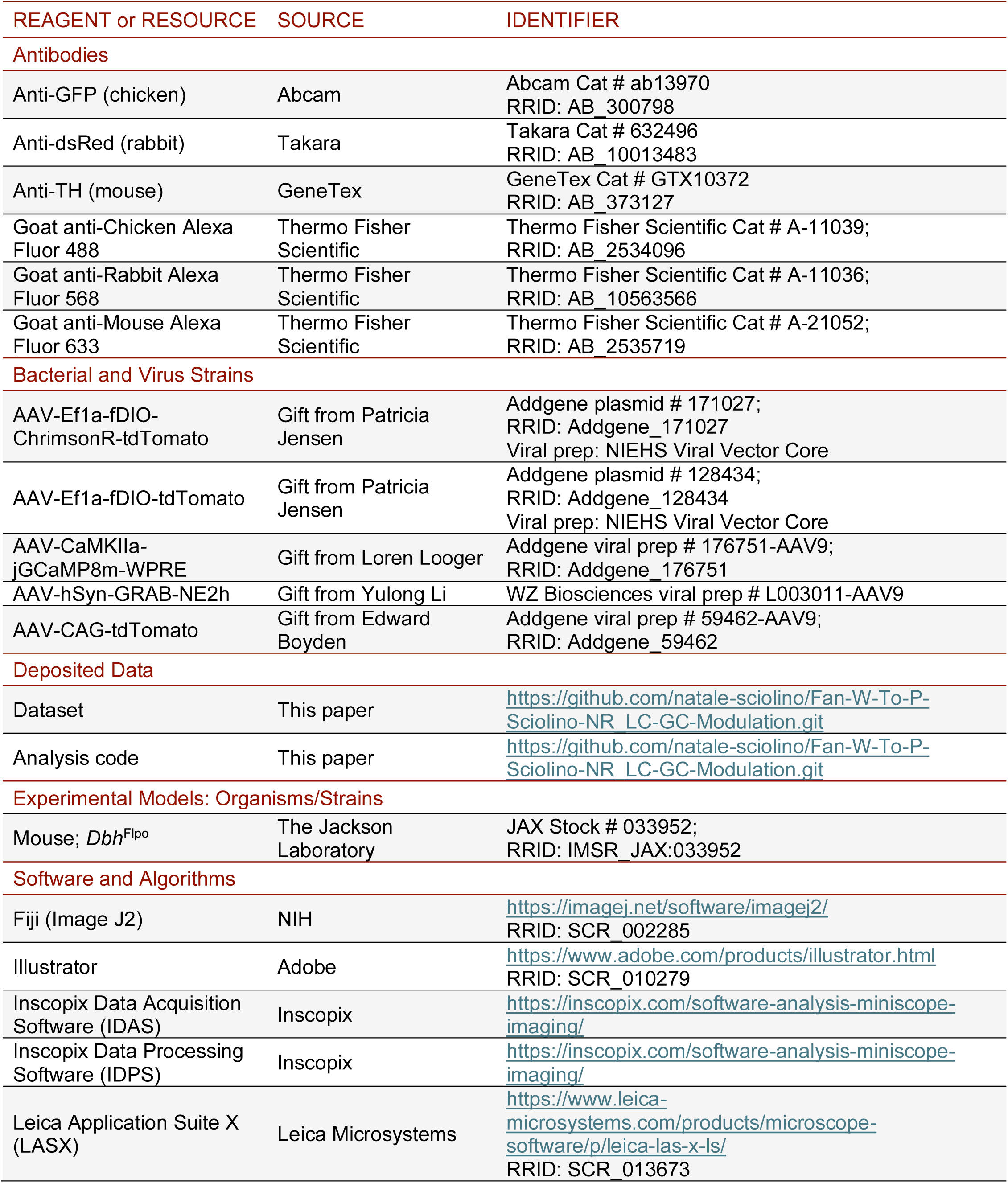
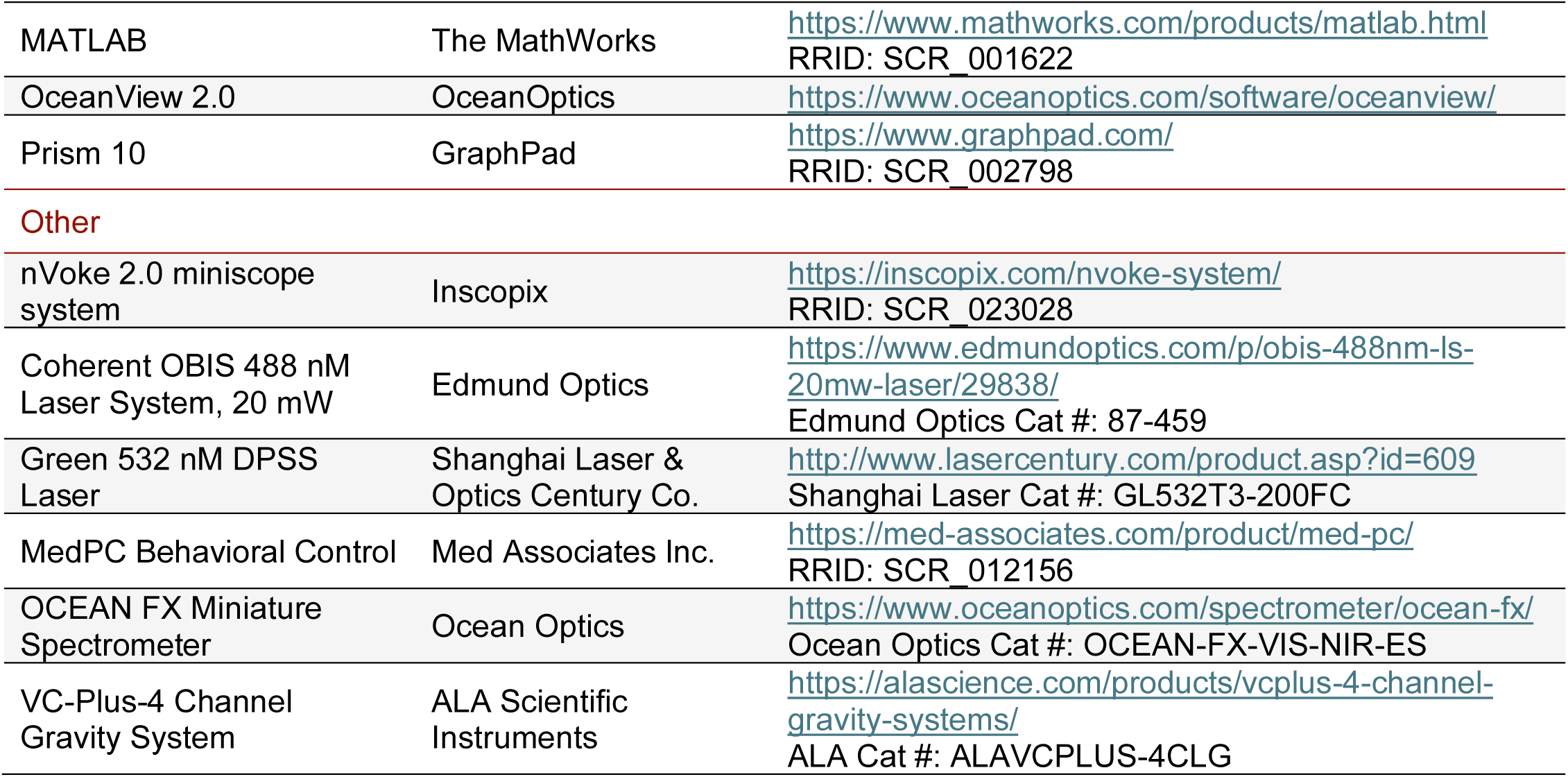

## RESOURCE AVAILABILITY

### Lead Contact

For questions or additional information, please contact the Lead Contact, Natale Sciolino. All reasonable requests will be accommodated.

### Materials Availability

No new or unique reagents were generated.

### Data and Code Availability

All neural and behavioral datasets, along with the original analysis code, have been deposited in Github and are publicly accessible as of the publication date. Corresponding DOIs are provided in the Key Resources Table. Additional information necessary to reproduce or reanalyze the findings described in this work is available from the Lead Contact upon request.

## EXPERIMENTAL MODEL AND SUBJECT DETAILS

Adult male and female mice older than postnatal day 60 (>P60) were used in all experiments. *Dbh*^Flpo^ mice^17^ were maintained as heterozygotes on a C57BL/6J background. Mice were housed under a reversed 12-hour light/dark cycle, with lights off at 9 A.M. (Zeitgeber Time 0, ZT0). All experiments occurred during the dark period. Mice had *ad libitum* access to food and water, except during experiments requiring water restriction. Prior to surgery, mice were housed in groups. All experiments were performed in accordance with the University of Connecticut’s Animal Care and Use Committee regulations and followed the *Guide for the Care and Use of Laboratory Animals*.

## METHOD DETAILS

### Survival Surgical Procedures

All survival surgical procedures were performed under 1-4% isoflurane anesthesia using aseptic techniques. Mice were positioned in a stereotaxic frame (Model 942, Kopf Instruments) atop a warming pad (RT-0514, Kent Scientific) to maintain body temperature. Anesthesia was delivered via a nose cone and continuously monitored throughout the procedure to ensure proper depth. For analgesia, ketoprofen (5 mg/kg i.p., Covetrus) was administered during surgery and again at 24- and 48-hours post-operatively for all procedures.

### Intracranial Viral Injections

Viral constructs were packaged in the indicated serotypes, and titers are reported as genome copies per milliliter (GC/mL) following dilution in physiological saline. Intracranial injections were performed after a midline scalp incision, followed by skull exposure and leveling. Unilateral craniotomies were made using a surgical drill (OmniDrill 35, WPI). Viral solutions were loaded into glass micropipettes pulled from capillary tubes (Fisher Scientific, Part 21171), backfilled with mineral oil (Fisher Scientific, Part O1211), and connected to a Nanoliter 2020 injector (WPI).

For miniscope imaging with LC activation, *Dbh*^Flpo^ mice received unilateral LC injections of AAVs encoding either the Flp-dependent optogenetic activator ChrimsonR^45,105^ (AAV-Ef1a-fDIO-ChrimsonR-tdTomato; serotype 5; 5 × 10^12^ GC/mL; Lot #: 09012021; *n* = 13 mice) or a fluorescent control^45^ (AAV-Ef1a-fDIO-tdTomato; serotype 5; 2.9 × 10^12^ GC/mL; Lot #: 09012021; *n* = 3 mice). In the same surgery, a CaMKIIa-driven calcium indicator (GCaMP8m) was injected in the ipsilateral GC using AAV-CaMKIIa-jGCaMP8m-WPRE (serotype 9; 1.15 x 10^13^ GC/mL; Lot #: V197511) to target pyramidal neurons. For LC injections, 500 nL was delivered at 100 nL/min. Coordinates, relative to bregma were: anterior-posterior (AP) -5.45 mm, medial-lateral (ML) -1.25 mm, and dorsal-ventral (DV) -3.85. We used unilateral LC manipulation because LC projections to cortex are predominately ipsilateral (∼95%)^106^. For GC injection, 500 nL was delivered at 50 nL/min. Coordinates, relative to bregma: AP +1.2 mm, ML -3.3 mm, DV -1.95 (from brain surface).

To monitor NE dynamics in the GC during LC activation, a separate cohort of *Dbh*^Flpo^ mice (*n* = 3 mice) received unilateral AAV injections of Flp-dependent ChrimsonR^45,105^ into the LC and a 4:1 cocktail of AAVs expressing the constitutive NE biosensor GRAB_NE2h_^107^ (AAV-hSyn-GRAB-NE2h; serotype 9; titer: 8 x 10^12^ GC/ mL; Lot #: 20240529) and tdTomato (AAV-CAG-tdTomato; serotype 9; titer: 4.0 x 10^10^ GC/mL; Lot #: V197427) into the ipsilateral GC. For this GC injection, 500 nL of the viral cocktail was infused at 100 nL/min. Coordinates relative to bregma: AP +1.2 mm, ML -3.3 mm, DV -2.0 (from brain surface).

After each injection, the micropipette was left in place for 5 minutes to minimize backflow. Incisions were sutured, and mice recovered on a heating pad for at least 24 h.

### Intracranial Optical Fiber and GRIN Lens Implantation

All procedures preceding viral injection, including pre-operative care, were conducted as previously described. This section outlines the implantation of custom fiber-optic probes and gradient refractive index (GRIN) lenses. Fiber-optic probes were constructed using 200 μm, 0.39 NA multimode fiber (FT200EMT, Thorlabs) housed in ceramic ferrules (MM-CON2007-2300, Precision Fiber Products) and were used for optogenetic stimulation in the LC or NE fluorescence detection in the GC. GRIN lenses (ProView Integrated Lens, 1 mm Ø x 4 mm length, with baseplate; Inscopix) were used for GC imaging.

For mice expressing either ChrimsonR or tdTomato in the LC, along with GCaMP8m in GC, a second surgery was performed ≥ three weeks after viral injection to implant the optical components. A small craniotomy was made above the LC and a larger one above the GC using a 1.4 mm inverted-cone bur (Panther). An optical fiber was unilaterally implanted above the LC at coordinates AP: -5.45 mm, ML: -1.25 mm, DV: -3.85 mm relative to bregma, and secured to the skull using Metabond (S371, S398, and S396, Parkell), while leaving the GC craniotomy open.

To guide GRIN lens placement above the GC, a flattened needle was lowered to 1.65 mm below the brain surface and then withdrawn. The GRIN lens was then implanted at 1.80 mm depth and secured with Metabond. For mice expressing ChrimsonR in LC and GRAB_NE2h_ in GC, a second surgery was performed three weeks post-injection to implant optical fibers above both regions. The LC fiber was implanted at AP -5.45 mm, ML -1.25 mm, DV -3.85 mm (bregma-referenced), and GC at AP +1.2 mm, ML -3.3 mm, DV: -2.0 mm (relative to the brain surface). Incisions were sutured, and post-operative care followed the protocol described above.

### Intraoral Cannulae (IOC) Implantation

A third surgery was performed to implant IOC in mice for miniscope recordings, at least two weeks after optical implantations. Each IOC consisted of a 1.5 cm segment of plastic tubing (1883T3, McMaster-Carr), with one end flared to secure a trimmed plastic washer (93785A812, McMaster-Carr). Mice were anesthetized via subcutaneous (S.C.) injection of ketamine (100 mg/kg) and xylazine (12.5 mg/kg) mixture (Covetrus).

A scalp incision was made to expose the lateral edge of the skull. A needle was attached to the non-flared end of the IOC, which was inserted lateral to the second/third maxillary molar, passed through the masseter muscle and beneath the zygomatic arch, and exited at the scalp edge behind the eye. After removing the needle, the exposed end of the IOC was connected to a push-in fitting (QSM-M3-3, Festo). Mice were placed in a stereotaxic frame equipped with a heating pad and maintained on 1-4% isoflurane anesthesia. The IOC was secured to the skull using Metabond and dental acrylic, and the incision was sutured closed. A fishing line plug was inserted into the IOC to prevent obstruction. To prevent infection, Penicillin G procaine (3,000 units/lb., Norbrook; 0.01 to 0.02 mL; intramuscular) was administered immediately post-surgery and again 48 hours later. All other post-operative care followed the protocol described above.

### Water Restriction Protocol

Following IOC surgery, mice were initially provided with water *ad libitum*. Starting at least two weeks post-surgery, they were gradually transitioned to a 23-hour water restriction protocol to stabilize fluid intake prior to neural recordings. Acclimation occurred over three days, with restriction periods progressively increasing from 12 to 23 hours. Once acclimated, mice received a minimum of one hour of water access daily, after passive tastant delivery (described below), to account for potential post-experiment satiety. Body weight and general health were monitored and recorded throughout restriction, which was limited to a maximum of two weeks.

### Passive Tastant Delivery

Taste stimuli were delivered using a gravity-driven, valve-controlled perfusion system (VC3-4PG, ALA Scientific Instrument). This system included four fluid reservoirs, each connected via separate 1/16″ PVC tubing (MMF-4, ALA Scientific Instruments) that converged at a custom-made manifold composed of four jointed miniature polyimide tubes (085-III, MicroLumen). Before each experiment, the IOC plug was removed, and the cannulae was cleaned using a fishing line to dislodge debris, followed by rinsing with deionized water-filled syringe, and flushing with compressed air to prevent obstruction.

To ensure consistent tastant delivery, mice were briefly anesthetized with 4% isoflurane, and any food debris or scar tissue near the IOC opening was removed with forceps. Mice were then placed in an empty arena and allowed to recover for at least 20 minutes. Immediately before each experimental session, the manifold was inserted into the IOC and secured using the push-in fitting. The manifold was designed so that its tip aligned flush with the IOC opening.

Three sets of taste stimuli were tested in separate experiments, including: 1) *Basic tastes*: 200 mM sucrose, 50 mM NaCl, 20 mM citric acid, and 1mM quinine, 2) *Sucrose concentrations*: 50 mM, 100 mM, 250 mM, and 500 mM, and 3) *Sucrose-NaCl mixtures*: 100 mM sucrose and 100 mM NaCl mixed at ratios of 90:10, 65:35, 35:65, and 10:90. Each experiment consisted of 96 trials with tastants presented in pseudo-random order with intertrial intervals of 25 to 35 seconds. During each trial, a solenoid valve opened for 0.08 seconds to deliver 8 µL of tastant. The perfusion system was rinsed and calibrated daily to ensure consistent fluid delivery. We did not include water rinses between tastant deliveries, consistent with prior studies^39,41,55,78^ because awake mice rapidly clear tastants rapidly through salivation. Any residual cross-trial tastant interactions would be expected to increase trial-to-trial variability rather than introduce systematic bias across conditions.

### *In Vivo* Miniscope Calcium Imaging

To monitor large-scale, single-cell Ca^++^ activity in the GC of freely moving mice, we used the nVoke 2.0 miniature microscope system (Inscopix). The miniscope was secured to the baseplate by tightening a small set screw and connected to a commutator (Inscopix), which enabled high-speed video and signal transmission while minimizing cable entanglement. The commutator was linked to a data acquisition (DAQ) box (Inscopix).

On the day of the experiment, mice were placed in an operant chamber (ENV-370W, Med Associates) within a sound-attenuating cubicle (ENV-017M, Med Associates) and given 15 minutes to habituate. The miniscope data were collected through the Inscopix Data Acquisition Software (IDAS). The miniscope has an excitation blue LED (455 ± 8 nm) that was set to 0.2 arbitrary unit power. The imaging gain (ranging from 2.0 to 3.0) was adjusted based on the fluorescence intensity histogram to ensure that the highest possible dynamic range was achieved without signal saturation. Data were collected at 20 frames per second (fps) with ∼50 ms exposure time. Synchronization of calcium imaging with tastant delivery and optogenetic stimulation was achieved using MedPC scripts (Med Associates) and transistor–transistor logic (TTL) signals delivered at 20 Hz.

### Optogenetic Stimulation Procedures

To activate LC neurons expressing ChrimsonR, a 532 nm green laser (Shanghai Laser & Optics Century Co., Ltd., China) was coupled to an optical patch cable (M72L02, M83L01, Thorlabs) and connected to the implantable probe via a mating sleeve (ADAL4-5, Thorlabs), which was wrapped in tinfoil to prevent light leakage. Laser power was adjusted to 8 mW at the fiber tip.

Two photostimulation paradigms were employed. In the *phasic stimulation* paradigm, half of the tastant delivery events were preceded by a 0.5-second train of laser pulses (20 Hz, 5 ms pulse duration). Tastants were administered 0.5 seconds after the photostimulation train ended. In the *tonic stimulation* paradigm, a total of 96 tastant delivery events were organized into eight blocks. Photostimulation at 5 Hz was applied during four of these blocks in the following the sequence: on, off, off, on, on, off, off, on. Within each stimulation block, the tonic pulse train began 45–55 seconds before the first tastant delivery and continued until 10 seconds after the final delivery. The subsequent block began 145–155 s after the preceding stimulation ended.

### Experimental Timeline

Two weeks following IOC implantation, mice were subjected to 23-hour water restriction to enhance motivation for tastant sampling. Prior to recording sessions, mice were habituated to wearing the miniscope during passive taste delivery in the testing chamber for three days. Each habituation session consisted of 96 tastant trials, including 200 mM sucrose, 50 mM NaCl, 20 mM citric acid, and 1mM quinine, allowing mice to become familiar with the stimuli.

The experimental design combined three sets of taste stimuli with two optogenetic stimulation paradigms, resulting in six distinct conditions: 1) Basic tastes–phasic stimulation, 2) Sucrose concentrations–phasic stimulation, 3) Mixtures–phasic stimulation, 4) Basic tastes–tonic stimulation, 5) Sucrose concentrations–tonic stimulation, and 6) Mixtures–tonic stimulation. These conditions were tested in the order listed above. Experimental sessions were occasionally separated by rest days, during which mice received 1 hour of water access without undergoing testing.

### Fiber Photometry Recordings of Photostimulation-Evoked NE Release

Five weeks post-optical fiber implantations, *in vivo* fiber photometry recordings were conducted to measure NE release in the GC following optogenetic stimulation of LC neurons. Recordings were conducted using a custom-built spectrally resolved fiber photometry system^45,48,108^. Excitation light at 488 nm (OBIS 488LS-20, Coherent) was adjusted to ∼60 µW power and directed into a fluorescence cube (DFM1, Thorlabs), reflected by a dichroic mirror (ZT488/561rpc-UF1, Chroma), and focused via an achromatic fiber port (PAFA-X-4-A, Thorlabs) onto a patch cable (M83L01, Thorlabs). The distal end was connected via a quick-release interconnect (ADAL3, Thorlabs) to an optical probe mounted on the mouse skull.

Fluorescence emitted from the tissue traveled back through the same optical path, passed through the dichroic mirror, and was filtered (ZET 488/561m, Chroma). The filtered signal was routed through a fiber port (PAF2S-11A, Thorlabs) and transmitted to a spectrometer (Ocean FX, Ocean Optics) via patch cable (M200L02-A, Thorlabs). Time-resolved fluorescence emission spectra of GRAB_NE2h_ and TdTomato signals were acquired using OceanView software (version 2.0.8) at 25 frames per second, with a 31 ms integration time. Synchronization of calcium imaging and optogenetic stimulation was achieved using MedPC (Med Associates) scripts and TTL signals delivered at 25 Hz.

Two recording sessions were conducted to test distinct photostimulation paradigms for activating ChrimsonR-expressing LC neurons and their effects on NE release in GC. In the first session, mice received *phasic stimulation,* consisting of 0.5-second trains of light pulses (5-ms pulse duration; 532 nm) delivered at 20 Hz. In the second session, mice received *tonic stimulation* at 5 Hz pulses. Both stimulation conditions used the same trial structure as the phasic and tonic miniscope experiments, respectively, except without tastant delivery.

### Brief-Access Taste Testing

Palatability of the four tastants used in the palatability experiment was assessed using a brief-access taste test. Eight *Dbh*^Flpo^ mice (separate from the above experimental subjects) were water restricted for 23 h prior to testing. During testing, mice licked a metal spout fitted with four microtubes, each connected to a reservoir containing one tastant solution. Licks were detected with a lickometer (Med Associates ENV-250), and each lick triggered delivery of ∼1 μL of tastant via the opening of a solenoid valve.

A trial began with the mouse’s first lick on the spout; the spout remained accessible for 10 s before being retracted. After a 10-s inter-trial interval, the spout was extended again to allow initiation of the next trial. Each session lasted 30 min, during which tastants were delivered in pseudo-random order and lick timestamps were recorded. The tastants tested—200 mM sucrose, 50 mM NaCl, 20 mM citric acid, and 1 mM quinine—were identical to those used in the palatability experiment.

### Tissue Collection and Processing

At the conclusion of the experiments, brain tissue was collected from all mice to verify viral expression and optical implant placement. Mice were deeply anesthetized with isoflurane and perfused transcardially with 1% phosphate-buffered saline (PBS) containing heparin (1:1000), followed by 4% paraformaldehyde in PBS (PFA/PBS), also supplemented with heparin. Brains were post-fixed overnight in 4% PFA/PBS at 4°C on a shaker, then cryoprotected in 30% sucrose in PBS for 48 hours. Following cryoprotection, brains were embedded in optimal cutting temperature (O.C.T.) compound (Thermo Fisher), frozen, and stored at −80°C. Coronal sections (40-µm) were cut using a cryostat (Leica CM 3050S) and transferred into a cryoprotectant storage solution containing 30% sucrose, 30% ethylene glycol, and 40% 0.1 M phosphate buffer. Sections were stored at −80°C until further processing.

### Immunohistochemistry

Brain sections were rinsed in 0.1 M PBS and blocked for 1 hour at room temperature on a shaker in 5% normal goat serum made in 0.1% Triton PBS. Sections were then incubated overnight at 4°C in primary antibodies, including chicken anti-GFP (1:10,000; Abcam; Lot # GR3361041-14), rabbit anti-DsRed (1:1000; Takara; Lot # 2103116), and mouse anti-TH (1:500; GeneTex; Lot # 822104279). The next day, tissues were washed in 0.1% Triton X-100 dissolved in 0.1 M PBS and incubated for 2 hours at room temperature on a shaker with secondary antibodies (1:1000; Fisher), including goat anti-chicken Alexa 488 (Lot # 2304258), goat anti-rabbit Alexa 568 (Lot # 2273773), and goat anti-mouse Alexa 633 (Lot # 2304276). After final rinses in PBS, brain sections were mounted onto slides and cover slipped using Prolong Diamond Anti-Fade mounting medium with DAPI (4′,6-diamidino-2-phenylindole; P36971, Thermo Fisher).

### Digital Image Processing

Immunofluorescent-labeled brain sections were imaged at 10x magnification using a Leica DM6 epifluorescent microscope (Leica Microsystems, Wetzlar, Germany). To enhance image clarity and reduce out-of-focus blur, all DM6 images were deconvolved using the Leica Instant Computational Clearing algorithm (LAS X 3.8.1). High-magnification images of the regions of interest were acquired at 40x using a Lecia SP8 spectral confocal microscope (Leica Microsystems). Digital images were exported from LASX software (v3.7.5) as uncompressed .lif or .tif files and opened in FIJI (v2.0) for further analysis. z-stacks were converted to maximum-intensity projections, and image adjustments were limited to uniform brightness and contrast changes to enhance fluorescent signal visibility.

### Data Presentation and Visualization

All graphical illustrations were made using Adobe Illustrator, Photoshop, GraphPad Prism, and MATLAB.

## QUANTIFICATION AND STATISTICAL ANALYSIS

Data analysis was performed using custom scripts written in MATLAB (MathWorks) and Prism 10 (GraphPad).

### Miniscope Movie Pre-Processing and Cell-Identification

Miniscope recordings were pre-processed using Inscopix Data Processing Software (IDPS), including spatial and temporal down-sampling, bandpass filtering, and motion correction. Cell bodies were identified using constrained non-negative matrix factorization (CNMFe)^109^, producing fluorescence traces in units of dF/noise. Extracted cells and traces were visually inspected to remove poorly identified cells. The traces were down-sampled to 10 Hz.

### Identification of Taste-Discriminative and LC-Modulated Neurons

All analyses were performed in MATLAB unless noted otherwise. Neuronal activity was aligned to tastant delivery and baseline-corrected by subtracting the mean response during the 1–4 s window preceding delivery. Aligned responses were pooled across subjects to construct pseudo-populations. Analyses focused on the 5 s post-tastant delivery period, divided into ten 500-ms time bins.

To identify stimulus-discriminative neurons, repeated-measures (RM) two-way ANOVAs (tastant × time bin) were performed on each neuron. Neurons were classified as stimulus-discriminative if either the tastant main effect or the tastant × time interaction was significant (*P*<0.05, two-tailed). To ensure robust evoked responses, we applied two additional criteria: (i) paired *t*-tests comparing each post-delivery time bin with baseline, retaining neurons with at least one significant time bin at *P*<0.001; and (ii) mean evoked response that differed from baseline by ≥1 dF. This procedure was applied separately to stim and no-stim trials, and neurons identified in either condition were combined. Thus, neurons were included in the stimulus-discriminative category if they met criteria in either condition, including cases near the classification boundary.

LC-modulated neurons were identified using RM two-way ANOVAs (stim-condition × time bin) separately for each tastant. Neurons were classified as LC-modulated if either the stim-condition main effect or interaction reached significance (*P*<0.01) for at least one tastant. The proportion of LC-modulated neurons in experimental and tdTomato control mice was compared using exact binomial tests (performed in R).

### Population Decoding

To decode tastant identity, linear discriminant classifiers (MATLAB *fitcdiscr* function) were trained on time-bin-resolved population responses. Decoding accuracy was evaluated via 6-fold cross-validation. To estimate performance for a given population size, 100 subpopulations were constructed by sampling neurons with replacement, and mean accuracy was computed. To test differences between stim and no-stim conditions, trial labels were permuted 1,000 times for each population size to generate a null distribution. The observed decoding difference was compared to the null distribution (two-tailed *P*-value). Significance across population sizes required at least three sizes with *P*<0.05. If this criterion was not met, no significance (n.s.) was noted on graphs.

### Response–Stimulus Correlation

Attribute-relevant information at the single-neuron level was quantified by computing Kendall’s rank correlation coefficient (Kendall’s tau) between each neuron’s mean responses to tastants and the ranked attribute values. LC stimulation effects were assessed by comparing the absolute values of tau across stim and no-stim conditions using RM two-way ANOVA (stimulation × time) followed by Bonferroni *post hoc* tests (using GraphPad Prism).

### Principal Component Analysis (PCA) and LC-Induced Transformations

Because PCA is sensitive to response magnitude, each neuron was normalized by its response to its preferred tastant. PCA was applied to mean population responses for each trial type (4 tastants × 2 stimulation conditions), using all baseline and 10-s post-delivery time points. This procedure stabilizes the PCA axes by fitting them to stimulus-averaged responses rather than noisy single-trial responses. Single-trial responses were averaged within the high response–stimulus correlation window (1.5–4.5 s post-delivery) and projected onto the PCA axes derived from trial-type means. Quantifications used the first six principal components (∼75% variance).

The palatability axis was defined as the vector connecting palatable (sucrose, NaCl) and aversive (citric acid, quinine) centroids, computed separately for stim and no-stim trials. Responses were projected onto this axis and normalized such that palatable centroid = 1 and aversive centroid = −1. Mixture-ratio and sucrose-concentration axes were defined similarly, except end-range tastants were used. Dynamic range was the difference between projection scores of the end-range tastants. The angle between stim- and no-stim-derived axes were computed using their dot product. LC-induced changes in tastant-pair separation were normalized by the no-stim dynamic range. We used normalized distance differences rather than distance ratios because ratios can become unstable when baseline distances are small.

Dynamic range changes, axis rotations, and pairwise distance changes were assessed by permuting trial labels between stim and no-stim condtions 1,000 times. Permutation-derived null distributions yielded two-tailed *p*-values for dynamic range ratios and one-tailed *p*-values for axis rotation. The latter choice assumed that LC-induced rotation would increase, rather than decrease, the angle that arose from variability in axis estimation. Pairwise distance changes were Bonferroni corrected for six comparisons. Dispersion orthogonal to the attribute axis was calculated as the mean perpendicular distance of points from the axis; LC-induced changes were assessed using the same permutation procedure. Procrustes transformations were used to align centroids between stim and no-stim conditions for visualization in the first three PCs.

For palatability, responses from both trial types were projected onto the no-stim-derived axis, and scores were compared using RM two-way ANOVA (tastant × stim condition) with Bonferroni-corrected *post hoc* tests (performed in GraphPad Prism).

### Modulation Type Analysis

Neuronal tuning was defined as mean responses to each tastant during 1.5–4.5 s post-delivery (same window as PCA). For stimulus-discriminative neurons, stim vs. no-stim responses were compared using three models: (i) *Null*: main effect of tastant only; (ii) *Scale–shift*: stimulation modifies tastant tuning via multiplicative and additive effects:

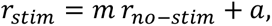

where 𝑟 denotes the tastant-evoked response, and 𝑚 and 𝑎 represent the multiplicative and additive factors, respectively; (iii) *Flexible*: tastant-specific changes are allowed by including interaction terms between tastant identity and stimulation condition, enabling arbitrary reshaping of the tuning profile. Null and flexible models were fit with linear least square regression (MATLAB *fitlm* function); scale–shift was fit with nonlinear least squares (*fitnlm*). Models were compared using Akaike Information Criterion (AIC) with hierarchical selection : scale–shift had to outperform null (ΔAIC > 2) to indicate modulation; flexible was selected only if it outperformed both. To prevent unstable nonlinear fits, scale–shift fits were constrained (m: 0–10; a: −10–10). Fits exceeding these bounds typically yielded extreme values and were thus excluded from the scale–shift category and evaluated only against the null and flexible models.

Proportions of scale–shift vs. flexible neurons were compared across experiments using logistic regression (binary response variable: modulation type; predictor: attribute), with pairwise Wald tests and Tukey correction (performed in R).

To relate multiplicative modulation to neuronal contributions to stimulus coding, we quantified each neuron’s contribution to attribute axes from the no-stim condition. Attribute axes, defined in PCA space, were projected back into neural activity space by multiplying each axis vector with the PCA coefficient matrix, yielding a vector of neuronal weights. Each neuron’s contribution was defined as the absolute value of its weight. For scale–shift modulated neurons, multiplicative factors were log-transformed to symmetrize expansive and compressive effects, and their relationship to neuronal contributions was assessed using Pearson’s correlation.

### Fiber Photometry Data Analysis

Raw photometry signals were preprocessed using a custom R script to linearly unmix the overlapping GRAB_NE2h_ and tdTomato signals, following established protocols^108,110^. To mitigate artifacts in GRAB_NE2h_ fluorescence signal (e.g., due to tissue displacement, cable bending during animal movement), the ratio of unmixed GRAB_NE2h_ to tdTomato signals (fluorescence ratio) was computed and used for all subsequent analyses, as performed previously^45,48,108^.

Fluorescence ratio traces were analyzed using custom MATLAB scripts. The percent change in fluorescence (%ΔF/F) was calculated by normalizing fluorescence ratios to their baseline mean values.

### Brief-Access Taste Test Analysis

Licking data from the brief-access test were first processed in MATLAB to calculate the mean lick counts for each 10-s trial. Lick counts were then analyzed using a RM one-way ANOVA, followed by Šídák’s *post hoc* comparisons (performed in GraphPad Prism).

---

**Figure S1.**
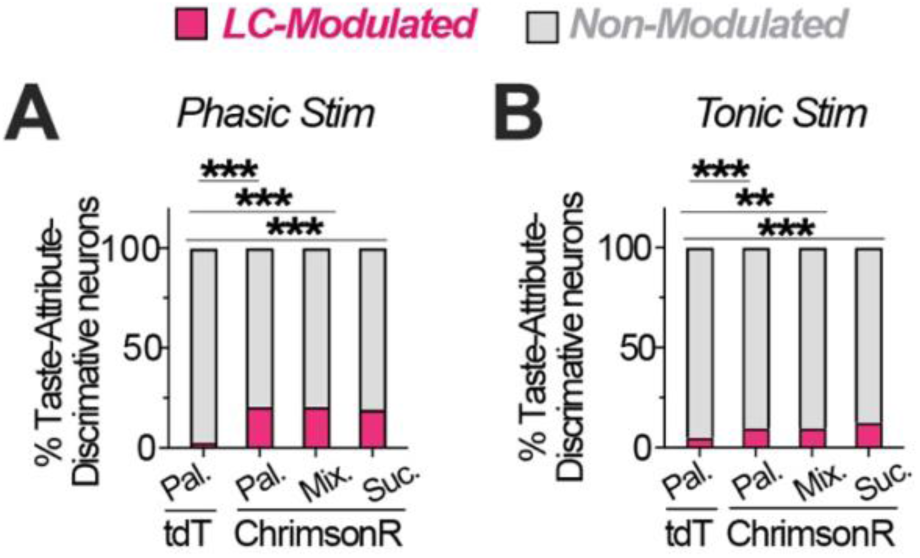
ChrimsonR Mice Have More GC neurons Identified as Modulated versus tdTomato Control. (A) Proportion of GC neurons identified as LC-modulated during phasic stimulation in tdTomato (tdT) control and ChrimsonR groups for palatability (Pal.), mixture ratio (Mix.), and sucrose concentration (Suc.) experiments. Across all attributes, ChrimsonR mice have more neurons modulated than tdT control (exact binomial tests): Palatability, 20.83% vs. 2.88% (\*\*\**P*<0.0001, 95% CI: 18.55–23.25%); Mixture ratio, 20.64% vs. 2.88% (\*\*\**P*<0.0001, 95% CI: 17.46–24.11%); Sucrose concentration, 19.26% vs. 2.88% (\*\*\**P*<0.0001, 95% CI: 15.94–22.94%). (B) Proportion of GC neurons classified as LC-modulated during tonic stimulation across taste attributes. ChrimsonR mice have more neurons modulated than tdT control (exact binomial tests): Palatability, 9.66% vs. 5.02% (\*\*\**P*<0.0001, 95% CI: 7.87–11.70%); Mixture ratio, 9.85% vs. 5.02% (\*\**P*<0.01, 95% CI: 6.59–14.01%); Sucrose concentration, 10.14% vs. 5.02% (\*\*\**P*<0.0001, 95% CI: 7.66–13.08%).

**Figure S2.**
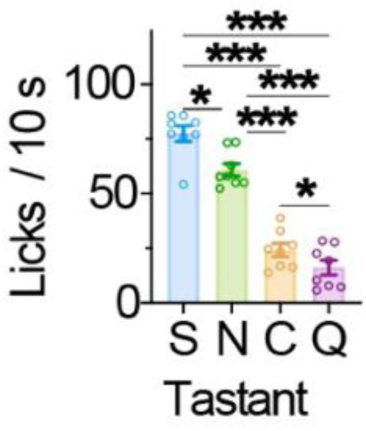
Licking Responses to Four Basic Tastes in the Brief-Access Test. Mean lick counts during 10-s, brief access to 200 mM sucrose (S), 50 mM salt (N), 20 mM citric acid (C), and 1 mM quinine (Q). A one-way RM ANOVA showed a significant tastant effect, *F*_1.538,10.76_*=* 126.7, *P*<0.0001. Šídák’s post-hoc tests confirmed the palatability order S > N > C > Q (*\*\*\*P*<0.001, \**P*<0.05). Data are mean ± SEM; dots represent individual mice (*n=*8).

**Figure S3.**
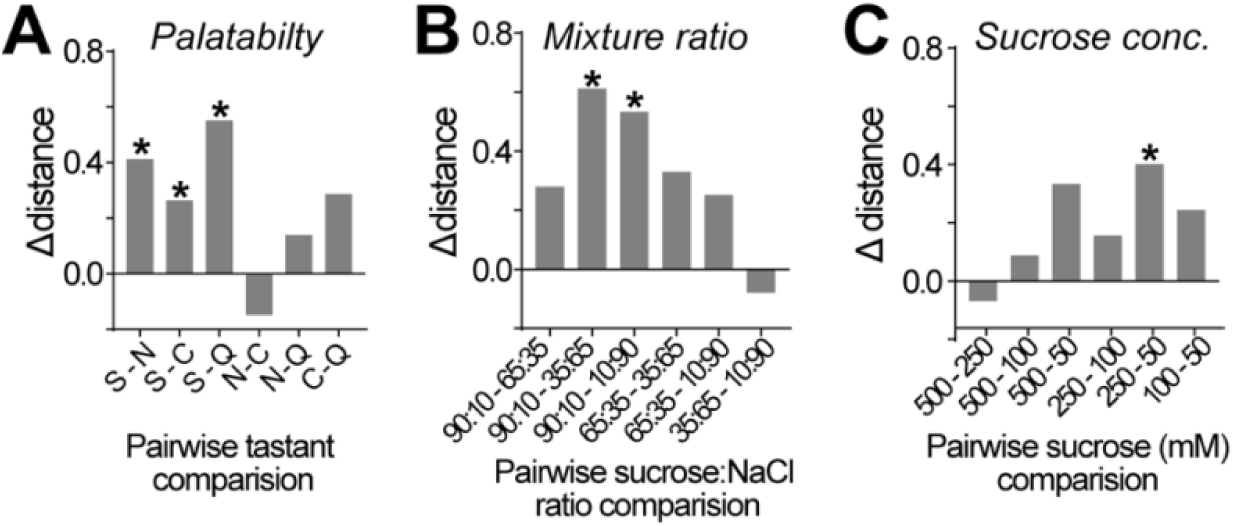
Phasic LC activation non-uniformly alters tastant separation along attribute-relevant axes. Changes in pairwise centroid separation (Δ distance) for (A) palatability, (B) mixture ratio, and (C) sucrose concentration. Δ distance was normalized to the no-stim dynamic range, with positive values indicate increased separation during phasic LC activation. Asterisks indicate significant permutation-test comparisons after Bonferroni correction (*p* < 0.05).

**Figure S4.**
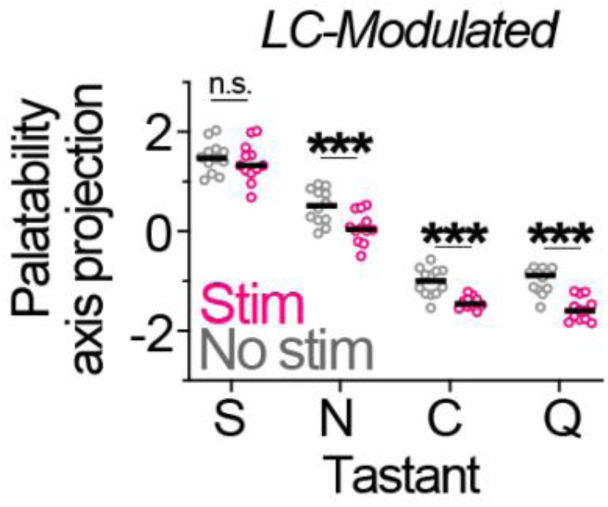
Phasic LC Activation Shifts Population Activity Along the Palatability Axis Towards a More Aversive Direction. Population responses of LC-modulated GC neurons under phasic LC stimulation and no-stimulation conditions were projected onto the no-stimulation-defined palatability axis. A two-way ANOVA revealed significant effects of Tastant (*F*_3,88_= 482.7, *P*<0.0001), Stimulation (*F*_1,88_= 40.22, *P*<0.0001), and Tastant x stimulation interaction (*F*_3,88_= 2.824, *P*=0.0433). Bonferroni’s post-hoc: *\*\*\*P*<0.001. n.s., non-significant. Black lines indicate means; dots represent individual trials (*n=*12 trials/condition).

**Figure S5.**
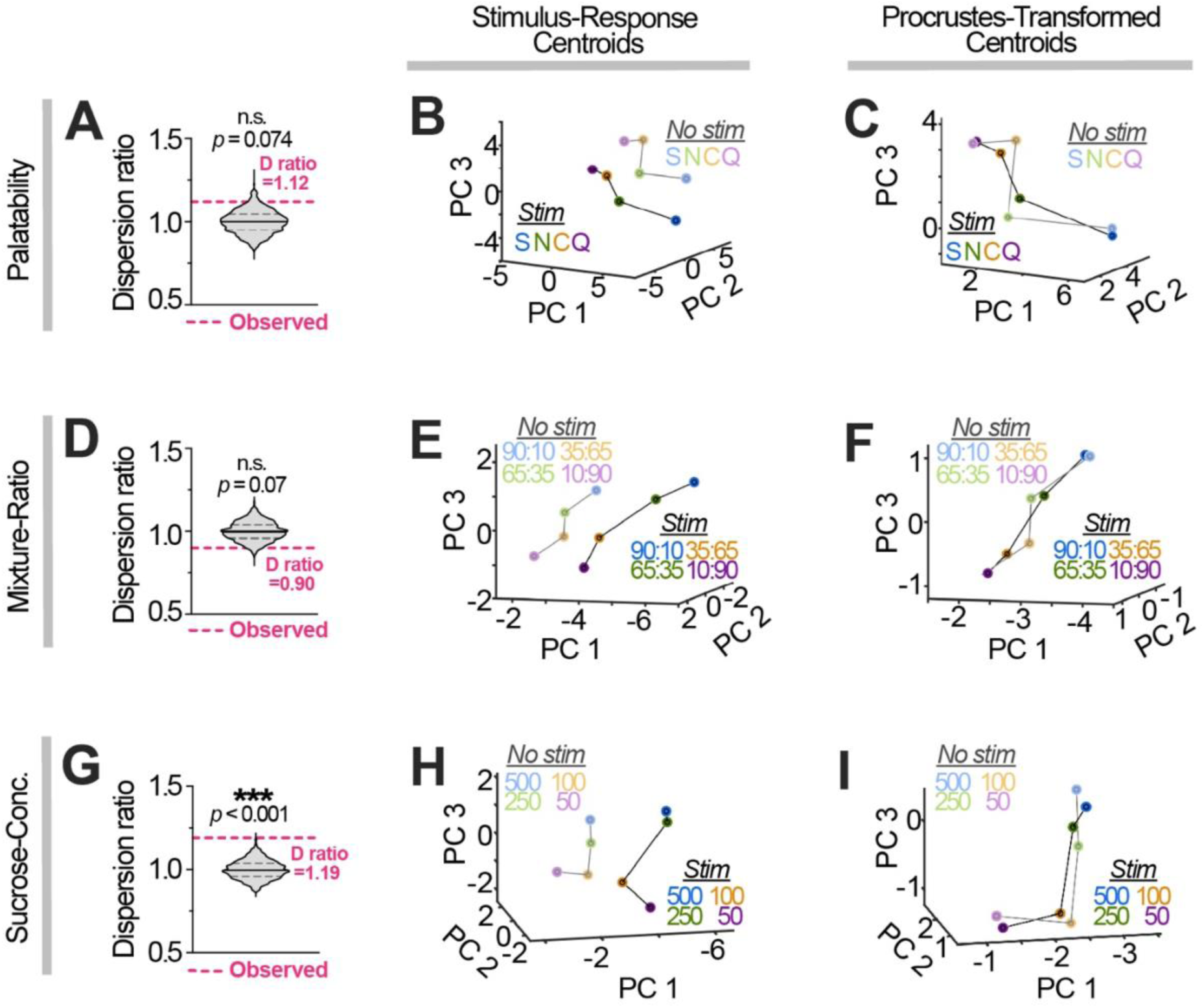
Effects of Phasic LC Activation on Tastant Responses in Directions Orthogonal to the Coding Axes in LC-Modulated GC Neurons. (A, D, G) Permutation tests for dispersion ratios for phasic LC-modulated GC neurons in the palatability (A), mixture-ratio (D), and sucrose-concentration (G) experiments. Dispersion is defined as the mean vertical distance of each taste response from the coding axis. The dispersion ratio compares stimulation vs. no-stimulation conditions. Violin plots show permuted distributions (median: solid black; quartiles dashed), with observed ratios in magenta. (B, E, H) Stimulus-response centroids for LC-modulated GC neurons in the palatability (B), mixture-ratio (E), and sucrose-concentration (H) experiments, projected into principal-component space with and without stimulation. Stimuli: palatability experiment—sucrose (S), NaCl (N), citric acid (C), and quinine (Q); mixture-ratio experiment—sucrose to salt ratios; sucrose-concentration experiment—listed concentrations (mM). (C, F, I) Stimulation centroids aligned to the no-stimulation centroids using Procrustes analysis for LC-modulated GC neurons in the palatability (C), mixture-ratio (F), and sucrose-concentration (I) experiments. Stimuli are as listed above.

**Figure S6.**
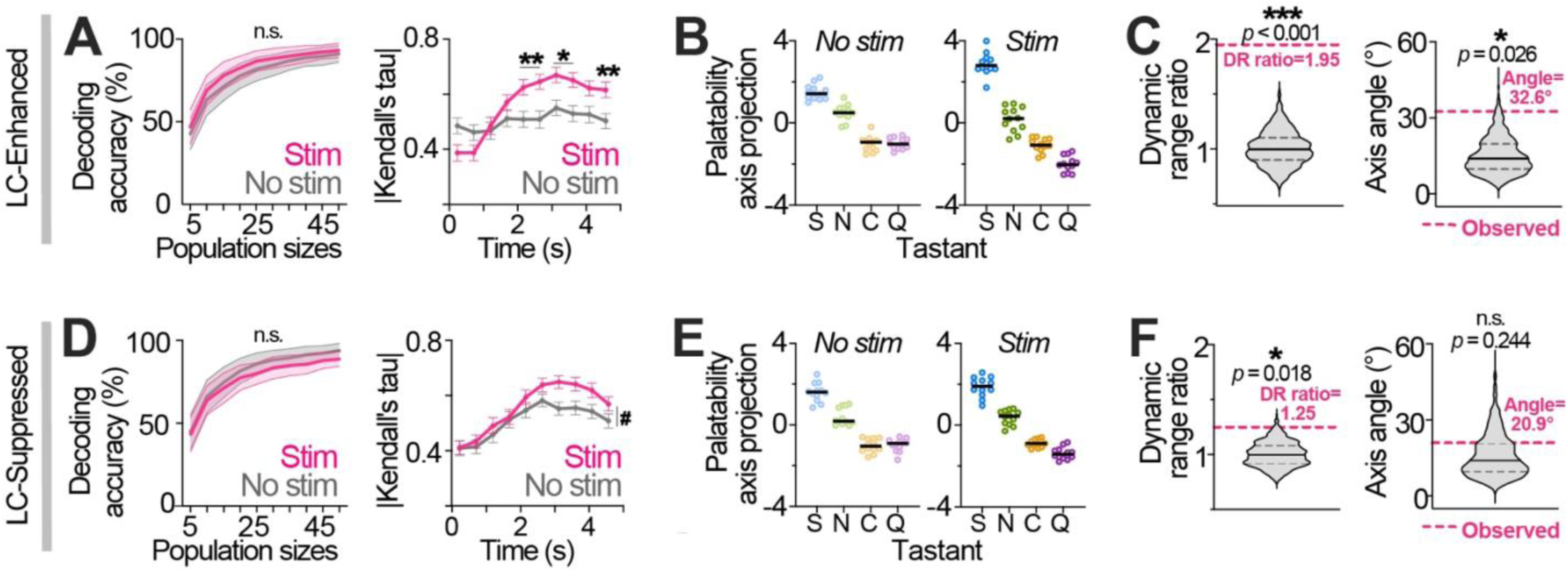
Modulation of Taste Encoding in GC Neurons Enhanced and Suppressed by Phasic LC Activation. (A) *Left.* Decoding accuracy for LC-Enhanced, taste-discriminative GC neurons across population sizes (mean ± SD of bootstrapped means). Permutation tests show no significant effects. *Right*. Correlations between response magnitude and palatability rank across time. Two-way RM ANOVA (time × stimulation): *F*_5.465, 606.6_= 7.044, *P*<0.0001. Main effects—stimulation: *F*_1.000, 111.0_= 8.064, *P*=0.0054; time: *F*_5.134, 569.9_= 13.19, *P*<0.0001. Bonferroni post-hoc: \*\**P*<0.01, \**P*<0.05. n.s., non-significant. Mean ± SEM (112 cells). (B) Population responses of LC-Enhanced neurons projected onto the respective palatability axis for stimulation (right) and no stimulation (left) trials. *n=*12 trials/condition. (C) Permutation tests for dynamic-range (DR) ratio (left) and palatability-axis angle (right). Violin plots show permuted distributions with medians (solid) and quartiles (dashed). Magenta lines indicate observed values. (D-F) Analyses for LC-Suppressed neurons follow the structure of panels A-C. (D) Decoding accuracy (left) and palatability correlations (right) for LC-Suppressed neurons. Permutation tests show no significant effects. Two-way RM ANOVA (time × stimulation): *F*_5.697,763.4_= 1.287, *P*=0.2630. Main effects—time: *F*_4.761, 637.9_= 21.47, *P*<0.0001; stimulation: *F*_1.000, 134.0_= 5.653, *P*=0.0188. ^#^*P*<0.05, stimulation. Mean ± SEM (135 cells). (E) Population responses projected onto the respective palatability axis for no-stimulation (left) and stimulation (right) trials. *n=*12 trials/condition. (F) Permutation tests for DR ratio (left) and palatability-axis angle (right) for LC-Suppressed neurons.

**Figure S7.**
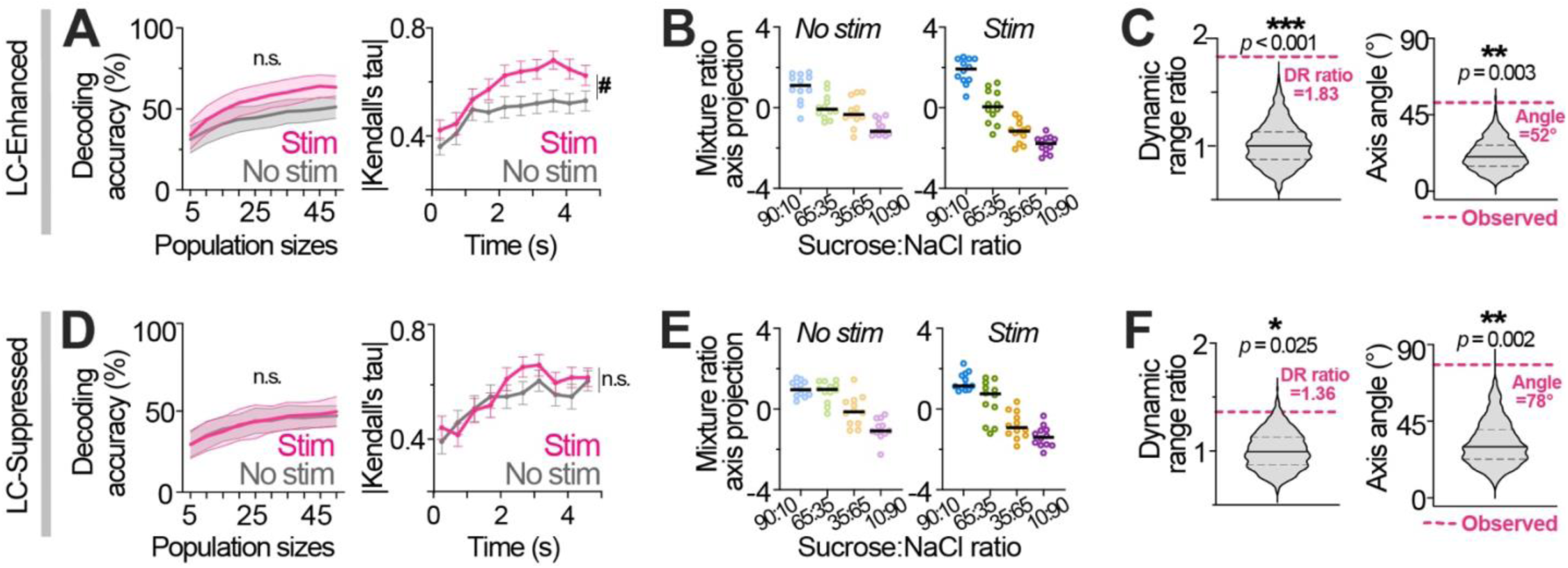
Modulation of Mixture-Ratio Encoding of GC Neurons Enhanced and Suppressed by Phasic LC Activation. (A) *Left.* Decoding accuracy for LC-Enhanced, mixture-ratio-discriminative GC neurons across population sizes (mean ± SD of bootstrapped means). Permutation tests show no significant effects. *Right*. Correlations between response magnitude and mixture-ratio rank across time bins. Two-way RM ANOVA (time × stimulation): *F*_5.347, 374.3_= 0.9743, *P*=0.4366. Main effects—time: *F*_4.657,326.0_= 11.09, *P*<0.0001; stimulation: *F*_1.000, 70.00_= 6.573, *P*=0.0125. ^#^*P*<0.05, stimulation. Mean ± SEM (71 cells). (B) Population responses of LC-Enhanced neurons projected onto the respective mixture-ratio axis for stimulation (right) and no stimulation (left) trials. *n=*12 trials/condition. (C) Permutation tests for dynamic-range (DR) ratio (left) and mixture-ratio-axis angle (right). Violin plots show permuted distributions with medians (solid) and quartiles (dashed). Magenta lines indicate observed values. (D-F) Analyses for LC-Suppressed neurons follow the structure of panels A-C. (D) Decoding accuracy (left) and mixture-ratio correlations (right) for LC-Suppressed neurons. Permutation tests show no significant effects. Two-way RM ANOVA (time × stimulation): *F*_4.497, 229.4_= 0.8674, *P*=0.4945. Main effects—time: *F*_5.756, 293.6_= 10.26, *P*<0.0001; stimulation: *F*_1.000, 51.00_= 0.5951, *P*=0.4440. n.s., non-significant main effect of stimulation. Mean ± SEM (52 cells). (E) Population responses projected onto the respective mixture-ratio axis for no-stimulation (left) and stimulation (right) trials. *n=*12 trials/condition. (F) Permutation tests for DR ratio (left) and mixture-ratio-axis angle (right) for LC-Suppressed neurons.

**Figure S8.**
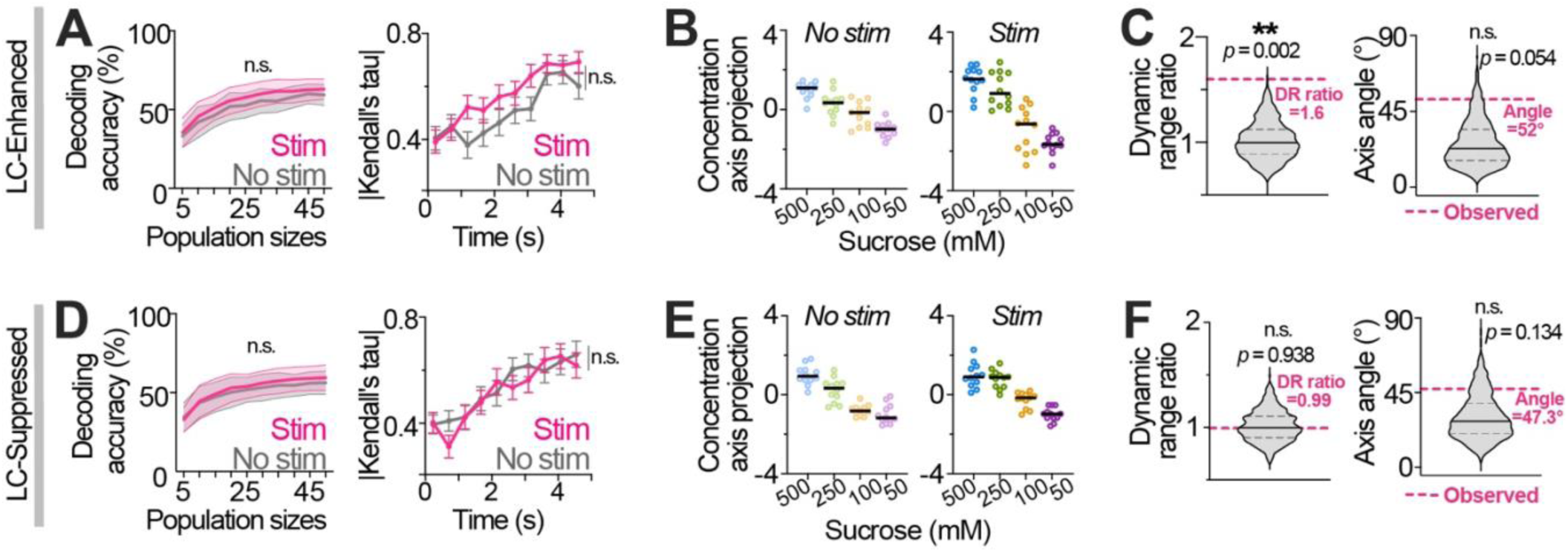
Modulation of Sucrose Intensity Encoding of GC Neurons Enhanced and Suppressed by Phasic LC Activation. (A) *Left.* Decoding accuracy for LC-Enhanced, sucrose-concentration-discriminative GC neurons across population sizes (mean ± SD of bootstrapped means). Permutation tests show no significant effects. *Right*. Correlations between response magnitude and sucrose-concentration rank across time. Two-way RM ANOVA (time × stimulation): *F*_5.452, 272.6_= 1.191, *P*=0.3126. Main effects—time: *F*_4.831, 241.5_= 13.23, *P*<0.0001; stimulation: *F*_1.000, 50.00_= 2.717, *P*=0.1056. n.s., non-significant stimulation effect. Mean ± SEM (51 cells). (B) Population responses of LC-Enhanced neurons projected onto the respective sucrose-concentration axis for stimulation (right) and no stimulation (left) trials. *n=*12 trials/condition. (C) Permutation tests for dynamic-range (DR) ratio (left) and concentration-axis angle (right). Violin plots show permuted distributions with medians (solid) and quartiles (dashed). Magenta lines show observed values. (D-F) Analyses for LC-Suppressed neurons follow the structure of panels A-C. (D) Decoding accuracy (left) and sucrose-concentration correlations (right) for LC-Suppressed neurons. Permutation tests show no significant effects. Two-way RM ANOVA (time × stimulation): *F*_4.800, 225.6_= 0.8637, *P*=0.5027. Main effects—time: *F*_4.320, 203.1_= 13.94, *P*<0.0001; stimulation: *F*_1.000, 47.00_= 0.1572, *P*=0.6936. n.s., non-significant main effect of stimulation. Mean ± SEM (48 cells). (E) Population responses projected onto the respective sucrose-concentration axes for no-stimulation (left) and stimulation (right) trials. *n=*12 trials/condition. (F) Permutation tests for DR ratio (left) and sucrose-concentration-axis angle (right) for LC-Suppressed neurons.

**Figure S9.**
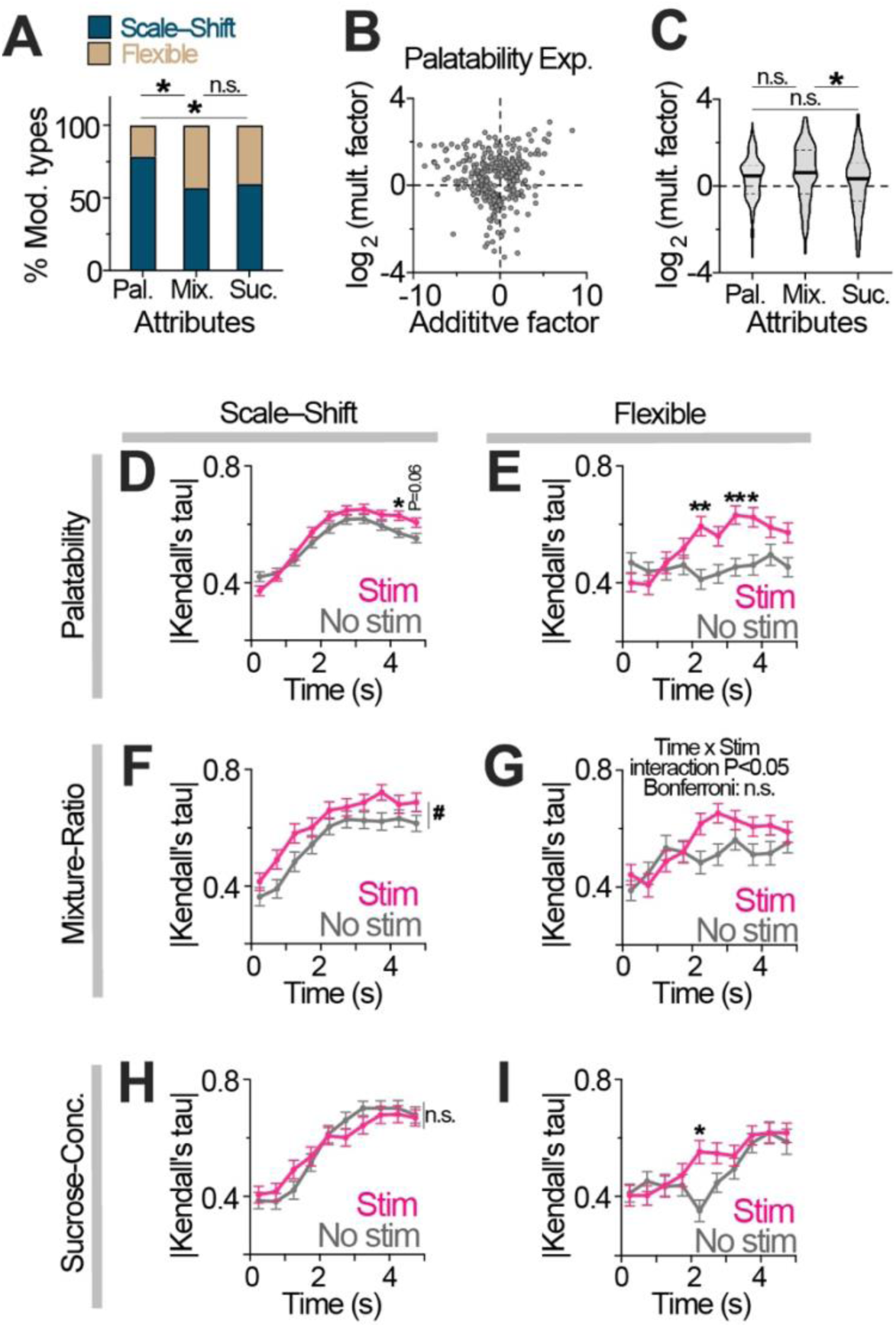
Additional Analysis of Scale–Shift and Flexible Modulation for Phasic LC Stimulation. (A) Proportion of GC neurons showing either scale–shift or flexible modulation during phasic LC stimulation for palatability (Pal.), mixture ratio (Mix.), and sucrose concentration (Suc.) experiments. Logistic regression revealed a significant effect of stimulus condition on the relative prevalence of scale-shift versus unconstrained modulation (likelihood-ratio test: *χ*²*_df=_*_2_= 37.80, *P*=6.21 × 10^-9^). Tukey-adjusted Wald tests showed that palatability stimuli had significantly higher odds of scale-shift modulation than both mixture (odds ratio = 2.8, *P*<0.0001) and sucrose stimuli (odds ratio = 2.5, *P*<0.0001), whereas mixture and sucrose conditions did not differ (odds ratio = 0.89, *P* = 0.86). (B) Additive factor versus log multiplicative factor for scale–shift-modulated GC neurons in the palatability experiment. (C) Log multiplicative factor distribution for scale–shift modulated GC neurons across taste attributes. Kruskal-Wallis: *H_3_*=6.716, *P*=0.0348. Dunn’s post-hoc: \**P*<0.05. n.s., non-significant. (D-I) Time-resolved correlations between neuron’s response magnitude and rank for palatability (D-E), mixture ratio (F-G), and sucrose concentration (H-I), shown separately for GC neurons exhibiting scale–shift (D, F, H) versus flexible (E, G, I) modulation during phasic LC stimulation. (D-E) Palatability correlations. Scale–shift neurons (D): Two-way RM ANOVA–Time × stimulation: *F*_6.240, 2034_= 3.225, *P*=0.0032; Time: *F*_4.924, 1605_= 66.54, *P*<0.0001; Stimulation: *F*_1.000, 326.0_= 4.037, *P*=0.0453. Flexible neurons (E): Time × stimulation: *F*_4.795, 417.2_= 5.090, *P*=0.0002; Time: *F*_4.630, 402.8_= 5.053, *P*=0.0003; Stimulation: *F*_1.000, 87.0_= 7.444, *P*=0.0077. Bonferroni post-hoc: \*\**P*<0.01, \**P*<0.05. Mean ± SEM (327 scale–shift cells, 88 flexible cells). (F-G) Mixture-ratio correlations. Scale–shift neurons (F): Two-way RM ANOVA–Time × stimulation: *F*_5.779, 554.8_= 0.4601, *P*=0.8215; Time: *F*_5.014, 481.4_= 30.27, *P*<0.0001; Stimulation: *F*_1.000, 96.0_= 6.588, *P*=0.0118. ^#^*P*<0.05, stimulation main effect. Flexible neurons (G): Time × stimulation: *F*_5.050, 363.6_= 2.422, *P*=0.0348; Time: *F*_5.662, 407.7_= 8.859, *P*<0.0001; Stimulation: *F*_1.000, 72.0_= 2.091, *P*=0.1525. n.s., non-significant. Mean ± SEM (97 scale–shift cells, 73 flexible cells). (H-I) Sucrose-concentration correlations. Scale–shift neurons (H): Two-way RM ANOVA–Time × stimulation: *F*_6.081, 681.1_= 1.639, *P*=0.1325; Time: *F*_4.992, 559.1_= 44.69, *P*<0.0001; Stimulation: *F*_1.000, 112.0_= 0.01902, *P*=0.8906. Flexible neurons (I): Time × stimulation: *F*_5.128, 384.6_= 2.719, *P*=0.0188; Time: *F*_5.204, 390.3_= 13.70, *P*<0.0001; Stimulation: *F*_1.000, 75.0_= 1.203, *P*=0.2762. Bonferroni’s post-hoc: \**P*<0.05. Mean ± SEM (113 scale–shift cells, 76 flexible cells).

**Figure S10.**
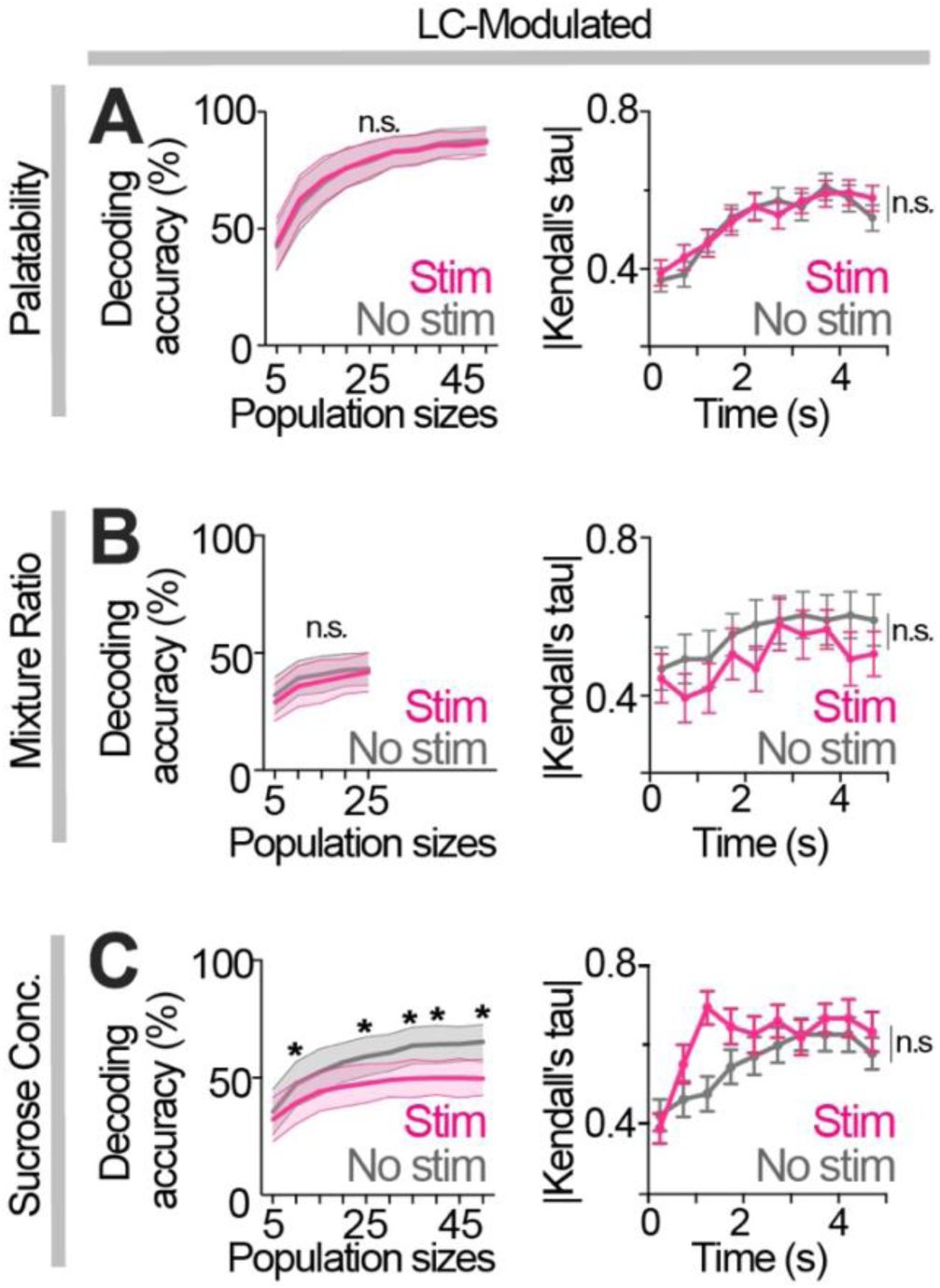
Effects of Prolonged Tonic LC Activation on Population Decoding and Stimuli Rank Correlations in LC-Modulated GC Neurons. (A-C) *Left.* Decoding accuracy across population sizes for tonic LC-modulated GC neurons in the palatability (A), mixture ratio (B), and sucrose concentration (C) experiments (mean ± SD of bootstrapped means). Permutation tests show no significant effects for palatability and mixture ratio experiments, whereas decoding accuracy was reduced in stim condition for the sucrose concentration experiment. \**P*<0.05. *Right*. Time-resolved correlations between each neuron’s response magnitude and stimuli ranking for each taste attribute. Palatability-discriminative neurons (A): Two-way RM ANOVA–Time × stimulation: *F*_5.072, 466.6_= 0.6119, *P*=0.6932; Time: *F*_4.769, 438.7_= 17.46, *P*<0.0001; Stimulation: *F*_1.000, 92.00_= 0.04783, *P*=0.8274. Mixture-ratio-discriminative neurons (B): Time × stimulation: *F*_3.758, 97.71_= 0.2440, *P*=0.9033; Time: *F*_4.604, 119.7_= 2.789, *P*=0.0235; Stimulation: *F*_1.000, 26.00_= 1.322, *P*=0.2607. Sucrose-concentration-discriminative neurons (C): Time × stimulation: *F*_5.910, 283.7_= 2.135, *P*=0.0505; Time: *F*_4.112, 197.4_= 7.428, *P*<0.0001; Stimulation: *F*_1.000, 48.00_= 2.533, *P*=0.1180. n.s., non-significant stimulation effects. Mean ± SEM (93 palatability cells; 27 mixture-ratio cells; 49 sucrose-concentration cells).

**Figure S11.**
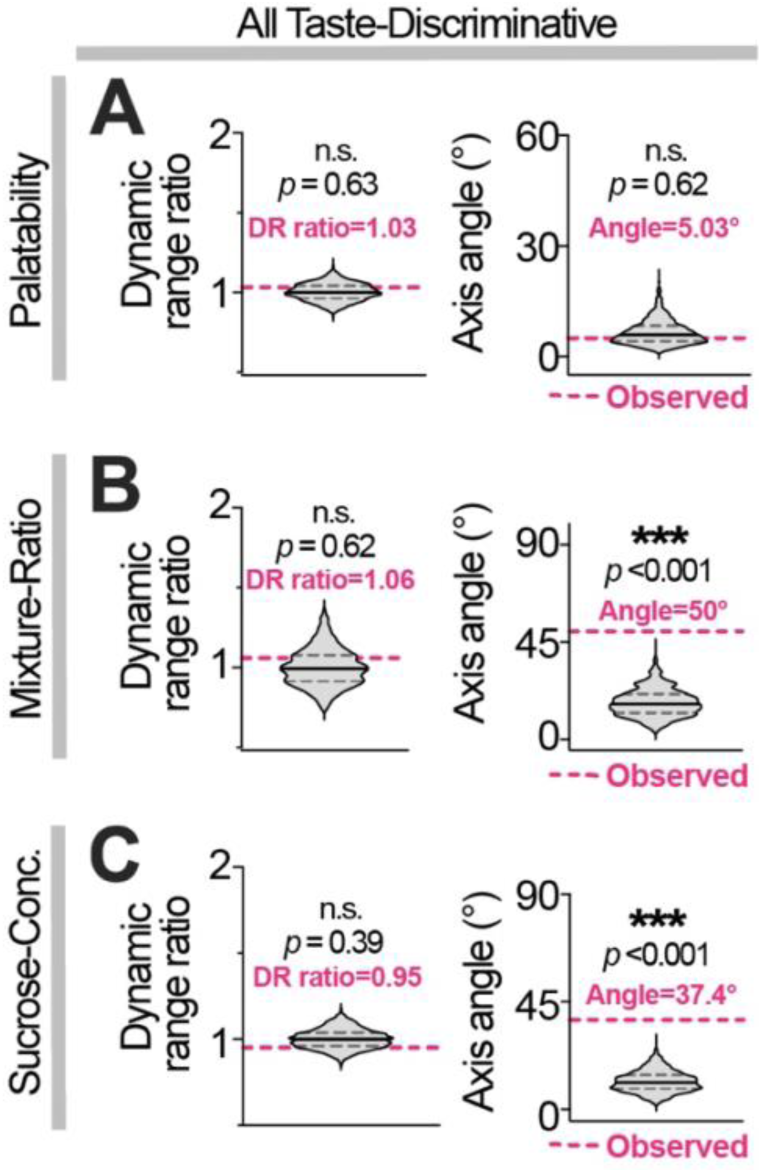
Prolonged Tonic LC Activation Had No Axis Stretching Effects and Rotated Palatability and Mixture-Ratio Axes. (A-C) Permutation tests for dynamic-range (DR) ratios (left) and axis angles (right) are shown for all stimulus-discriminative GC neurons in the palatability (A), mixture ratio (B), and sucrose concentration (C) experiments during tonic LC stimulation. Violin plots show permuted distributions (median: solid black; quartiles dashed), with observed values shown in magenta.

